# Precipitation and Extraction Methods for Protein Purification: A Meta-Analysis of Purification Performance and Cost-Effectiveness

**DOI:** 10.1101/2023.12.14.571684

**Authors:** John S. Decker, Utsuki Yano, Romel Menacho Melgar, Michael D. Lynch

## Abstract

For protein drug purification, packed-bed chromatography often remains both the predominant method and a bottleneck for cost and scalability. Accordingly, extensive efforts have been made to develop alternatives, such as precipitation and liquid-liquid extraction. Despite decades of development, such methods have been slow to see adoption in commercial processes. To diagnose the key barriers to implementation and guide future work, we have systematically reviewed studies of protein precipitation and liquid-liquid extraction. We classify the products, methods, and results of 168 publications representing 290 unique purification operations and analyze these operations in terms of both process economics and purification performance. Whereas it is generally assumed that precipitation and extraction methods will have lower costs than chromatography, we find that this is only the case under specific process conditions such as at a large manufacturing scale and low initial sample purity. Furthermore, we find that only a small number of the many precipitation and extraction methods reported to date have shown readiness for implementation in protein drug purification processes. Finally, we identify key factors governing both the economic and purification performance of this class of methods: first, that operating costs are almost entirely predictable by the ratio between the mass of phase-forming materials used and the mass of product protein yielded; second, that use of modern optimization techniques such as Design of Experiments is associated with significantly better purification performance and cost-effectiveness.

**Highlights:**

- Alternative separation purification methods are not always cheaper than chromatography
- The use of a combination of phase separating agents remains largely underexplored/underutilized
- Lower initial purity and increasing production scale favor phase-separation over chromatography
- The direct material usage rate is an important predictor of alternative separation cost-effectiveness
- Current alternative separation method development has largely ignored optimization of direct material usage rate

## 1. Introduction

The total market for recombinant proteins is currently estimated to be more than $270 billion, with a projected value of more than $400 billion by 2026 (BCC Market Research, 11/2021, 08/2021, 09/2021). More than 96% of these figures are accounted for by protein therapeutics, which are set apart from other applications by their stringent purity requirements and the associated high costs of manufacturing. For such drugs, manufacturing costs typically represent approximately 20% of total sales (Decker et al., 2020) and downstream processing typically represents about 75% of total manufacturing costs (Straathof, 2011). Therefore, protein purification processes alone cost more than $40 billion per year. The linchpin of these processes is undoubtedly conventional packed-bed chromatography (Kelley, 2017; Rathore et al., 2018), which is often also considered their major cost driver and productivity bottleneck (Farid, 2017; Gagnon, 2012).

Despite its limitations, conventional chromatography has remained the preeminent purification operation in biotechnology largely because of three key advantages. First, it is versatile: generally, when developing a purification process for a new protein product, almost all the necessary separations can be performed using some sequence of chromatography steps. Second, it is predictable: most resins in common use have a single dominant selectivity (e.g., electrostatic interactions or hydrophobicity), a relatively small number of easily-controlled variables drives the performance, and optimization strategies are well-established (Hibbert, 2012). Third, it is generally capable of high yield as well as high resolution even among proteins with similar properties. Any alternative separation operations seeking to displace conventional chromatography therefore face the imposing task of overcoming these advantages of versatility, predictability, selectivity and yield. Furthermore, they must do so in a cost-effective manner.

Nevertheless, in certain circumstances, alternative purification methods can dramatically lower process costs without sacrificing product quality, potentially enabling new markets or simpler and more flexible manufacturing processes (Decker et al., 2022, 2020). One highly active area of development for such alternatives deals with methods that achieve purification by differentially modulating protein solubility among two or more chemical phases (Dos Santos et al., 2017; Hekmat, 2015; Martinez et al., 2019; Roque et al., 2020; Soares et al., 2015). This class of methods can be referred to as phase separations or bulk separations (Przybycien et al., 2004) and includes precipitation and liquid-liquid extraction as among the most promising examples. These methods are generally considered attractive because of three potential advantages they offer over chromatography: they can typically achieve rapid separations (e.g., on the order of minutes), they require only simple and relatively inexpensive equipment, and they can be performed with inexpensive chemicals or reagents.

Despite their potential cost advantages, decades of development, and widespread use for non-therapeutic products and specific cases such as plasma proteins, phase separation methods have only seen slow and sparing uptake into commercial processes for recombinant protein therapeutics. Recent reviews identify only a handful of examples, including insulin, erythropoietin, and Apo2L (Dos Santos et al., 2017; Thömmes and Etzel, 2007). We set out to identify the reasons for this lack of implementation to date, and to guide future work in the field towards successful implementation.

Other recent reviews in the field have highlighted a relatively small number of successful examples of phase separation methods (Dos Santos et al., 2017; Hekmat, 2015; Martinez et al., 2019), surveyed the broad categories of such methods (Dos Santos et al., 2017; Roque et al., 2020), and addressed theoretical and engineering aspects of method development (Dos Santos et al., 2017; Hekmat, 2015; Martinez et al., 2019). However, prior reports have yet to systematically or comprehensively evaluate the literature in a way that enables identification of key factors separating success from failure in method development, nor have recent reports considered economic aspects of the methods in detail. They are therefore unable to diagnose the reasons for the low uptake of phase separation methods in protein drug processes or offer evidence-based suggestions for future work that are generalizable across diverse types of phase separation methods.

To address this gap, we present a meta-analysis of 168 publications reporting 290 distinct phase separation operations. First, we systematically survey the full range of purification performance for nearly 40 different phase separation methods, rather than only highlighting successes. Second, we identify methodological factors associated with increases in purification performance. Third, we conduct a detailed techno-economic analysis and identify the conditions under which phase separation methods are likely to be cost-effective compared to chromatography. Fourth, we assess the current implementation-readiness of various phase separation methods on the basis of both purification performance and economics. And finally, we suggest guidelines for future work in the field on the basis of this evidence.

## 2. Methods

### 2.1. Literature Inclusion Criteria

The scope of this analysis was limited to publications in scientific journals from 01/01/2000 to 01/19/2022 (the date of the literature search), in which the following elements could be identified: 1) a process for purifying a protein was reported; 2) a precipitation or aqueous extraction was performed; 3) a product yield was reported; 4) a degree of contaminant removal was reported; 5) original research was reported; and 6) full-text access was available in English and at the authors’ institution.

### 2.2. Literature Search

The PubMed database (National Library of Medicine (US)) was searched on 01/19/2022 to identify publications potentially matching the inclusion criteria (for the search term, refer to Supplemental Methods). 289 publications matching the inclusion criteria were identified, of which 168 were manually reviewed to achieve a representative sampling.

### 2.3. Literature Review

Each publication in the dataset was manually reviewed for annotation of key features describing the product, expression system, purification process, methodology, and results. Each publication was further divided into one or more individual records, where each record contained the information relevant to a distinct protein product and/or phase separation unit operation. For example, a publication reporting the use of two sequential precipitation operations to purify one product would comprise two records. In contrast, when a publication reported the evaluation of multiple phase separation unit operations in competition, rather than for sequential use as part of the same process, only the one showing the best results was recorded. From the 168 publications reviewed, the dataset for analysis consisted of 290 unique records of phase separation purification operations.

For each analyzed feature, when possible, data were transformed into standardized units and formats across all records. For a list and definitions of all features considered in this analysis, as well as approaches used to standardize data, refer to Supplementary Methods.

### 2.4. Classification of alternative separation methods

Information describing the reaction conditions of each purification operation was collected as completely as possible, including, as applicable, the concentrations and identities of each chemical or biological species present and the temperature, time, pH, and volumes involved. However, for the purposes of most analyses, methods were also more broadly grouped: first as using either precipitation or liquid-liquid extraction, and then by the types of agents used to effect the separation. Phase-forming agents were classified by general chemical properties (e.g., charge, size, polarity) and/or by the features of the product and contaminant proteins that they probe (e.g., affinity recognition or isoelectric point).

### 2.5. Classification of optimization methods

Records in the dataset were also classified in terms of the methods used to optimize the phase separation unit operation. Optimization methods were categorized in four ways. If nothing in the publication suggested whether or how variables were tested before selecting the final reaction conditions, we classified a study as having no optimization. If each variable was tested independently and final levels of each variable were chosen based on the independent optima, we classified a study as using one-factor-at-a-time optimization (OFAT). If variables were tested simultaneously and final levels of each variable were chosen based on the optimum from the multiple-variables tests, we classified a study as using multiple-factors-at-a-time (MFAT) optimization. Finally, if MFAT experimentation was supplemented by the use of statistical principles to select the tested variable levels, experimental order, or other features of the approach, we classified the study as using Design of Experiments (DoE).

### 2.6. Data Analysis, Visualization and Statistics

Data analysis, visualization, and statistics were performed in Python using standard methods via the libraries pandas (McKinney, 2010), seaborn (Waskom, 2021), Scipy (Virtanen et al., 2020), and Matplotlib (Hunter, May-June 2007). Visualization was also performed in GraphPad Prism (GraphPad Software, San Diego, CA).

Linear least-squares regressions were performed after taking the log_10_ transform of the x and y variables to linearize the data. Statistical differences in product yield were determined using non-parametric tests (the Kruskal-Wallis test and the Mann-Whitney U test for group and pairwise comparisons, respectively) because of the non-normality of the data. Differences in host cell protein removal were determined using one-way ANOVA for group tests and Welch’s t-test for unequal variances for pairwise tests. All multiple comparisons for pairwise tests were corrected by the Bonferroni method. Significance was determined at p < 0.05 or the appropriate corrected value.

### 2.7. Techno-economic Analysis

Techno-economic analyses were performed in Python using custom unit operation models for a generic phase separation protein purification method and for ion-exchange chromatography. Each operation was modeled to purify four amounts of product through the unit operation per year: 10 kg, 100 kg, 1000 kg, and 10000 kg. Whenever maximum equipment sizes were exceeded, the equipment was divided into as many equally-sized parallel units as necessary. Records from the dataset were included in the techno-economic analysis if they reported a measure of host cell protein removal, a measure of product yield, a product purity level prior to phase separation, and a description of the reaction conditions. For each record, these data were then used to parameterize models of both the phase separation operation as reported and an alternative ion-exchange chromatography operation processing the same input stream. The complete models may be viewed and tested as interactive Python scripts in a web browser at https://mybinder.org/v2/gh/jsd94/Decker-et-al-2022-Techno-economic-analysis/main. Additional details on techno-economic analyses can be found in Supplementary Methods.

## 3. Results

### 3.1. Dataset characterization

From 168 reports published between 2000 and January 2022, we identified a dataset of 290 distinct combinations of phase separation methods and product proteins (i.e., 290 data “records”). We began by surveying the existing use-cases of precipitation and liquid-liquid extraction for protein purification, focusing on four descriptors: the type of method used; the type of product purified; the type of host organism from which the product was purified (or, more generally, the source of the contaminant protein background); and the application for which the research was carried out.

We found that nearly 40 distinct types of phase separation method were in use. Of these, 15 were observed in 5 or more cases (Figure S1). However, while the five most common methods accounted for over half of all data, most methods reported were associated with only one or a few records. Therefore, most of the observed methods lack the level of replication or generalization to diverse products that would be desirable for making conclusions about their readiness for commercial implementation.

We also broadly classified products purified in the analyzed studies as monoclonal antibodies or antibody fragments, crude immunoglobulin fractions (Ig), enzymes, or none of the former (Figure S2A). While it would be ideal to further classify products based on structural features, the great variety of products in the dataset and the sparsity of sequence or structural information, made this classification unfeasible within the scope of this study. We additionally classified hosts as bacteria, mammalian tissue culture, fungi, plants, or animal-derived samples (e.g., serum), with over half of the data approximately equally divided between the first two categories (Figure S2B). In terms of research application, by far the most common was medical products, representing more than half of the data (Figure S2C). Additional sub characterizations of the dataset by these categories are available in Figures S3-7.

In the remainder of this analysis, we will interpret the data with a particular emphasis on using phase separation methods to purify medical products for three reasons. The first is that this scenario represents such a large fraction of the dataset. The second is that medical products make up by far the largest fraction of the overall recombinant protein market. The third is that this scenario is associated with the most stringent and most uniform purification requirements as well as the highest manufacturing costs, making the comparison to established alternative methods such as packed-bed chromatography both easiest and most meaningful.

To begin the analysis, we sought to extract information pertinent to the cost and the purification method description and performance (Table S1). However, with respect to performance variables, we found that in the literature on phase separation-based protein purifications, most were reported in 50% or less of cases (Figure S8). Furthermore, this sparsity of important data was not the result of a subset of highly incomplete records. Rather, almost all records in the dataset were substantially incomplete (Figure S9). This represents an important barrier to the progress of the field as well as to the uptake of these methods into commercial processes. For the field to progress, there needs to be more consistent reporting of the key metric required for a thorough evaluation. Nevertheless, sufficient information could be found to permit quantitative analyses of both purification performance and economics.

### 3.2. Purification performance of phase separation vs chromatography

Having generally described the contents of the dataset, we next turned to a quantitative evaluation of protein purification performance. The goal of this evaluation was to guide considerations of which phase separation-based methods should be considered for commercial adoption, and which require further study. To this end, we compared phase separation methods both to each other and to two of the most common chromatography-based methods in pharmaceutical bioprocessing, ion-exchange (IEX) and Protein A chromatography (ProA). For the purposes of comparison, we obtained data on IEX and ProA performance from the literature (see Tables S2-4).

The first aspect of purification performance we considered was removal of host cell proteins (HCPs), because no other measure of contaminant removal was widely reported in a manner that allowed direct comparison among different studies. HCP removal values (here shown as the log_10_ reduction value in absolute HCP mass, or LRV) could be identified for 21 different methods (Figure 1A). Of these, 14 were only represented by three or fewer datapoints. For the other 7 methods, the variation within methods was generally greater than the variation among methods. Thus, it was not clear that any one phase separation method or set of methods could be reliably predicted to perform better than another for an arbitrary new product.

**Figure 1:**
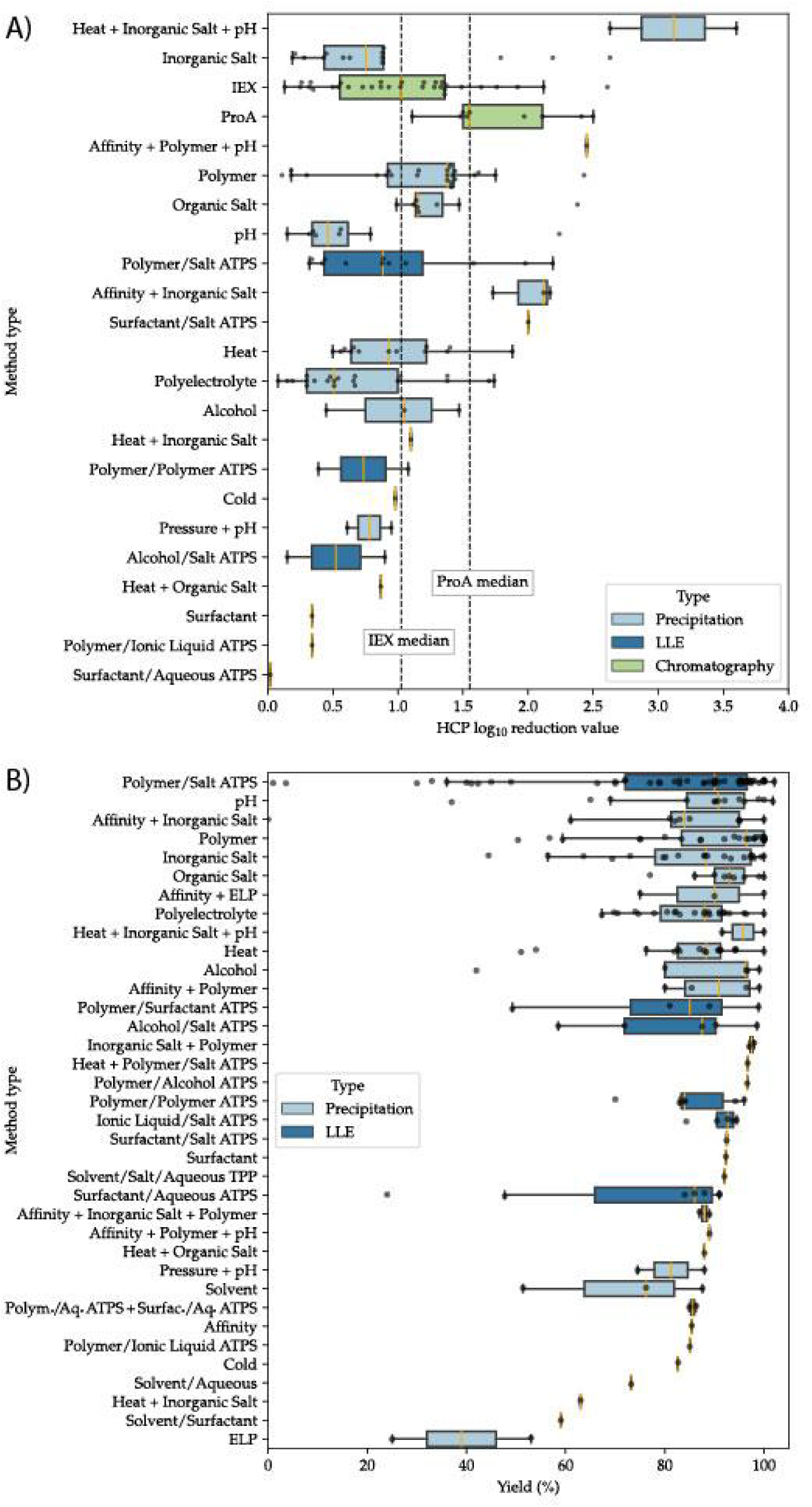
Performance of phase separation-based protein purification methods. (A) Host cell protein removal by method type for phase separation-based methods and two common types of chromatography. Methods are precipitations unless ending with ATPS (aqueous two-phase separation). (B) Product yields of phase separation-based protein purification methods showing many have demonstrated high product yields. LLE: liquid-liquid extraction. ATPS: aqueous two-phase separation. ELP: elastin-like polypeptide. TPP: three-phase partitioning. For both panels, box and whisker plots show the minimum, 25^th^ percentile, 75^th^ percentile and maximum values, respectively. Individual datapoints are shown as black circles. The median of each group is highlighted in orange.

When compared to chromatography-based alternatives, 11 and 14 phase separation methods had at least one report of HCP removal greater than the median value for ProA or IEX, respectively. However, median performance compared much less favorably. Among methods with two or more datapoints, only 2 and 5 methods had median HCP LRVs greater than the median for ProA and IEX, respectively. Moreover, 3 of these 5 methods were represented by only two or three datapoints each. Thus, while it is clear that phase separation-based methods can sometimes outperform chromatography in HCP removal, more work is required to validate the robustness and predictability of this performance under product or process variations.

Since monoclonal antibody (mAb) products are the most economically significant area of the reported research in the field, we conducted this evaluation separating mAb products (Figure S10A) from others (Figure S10B). In this case, while several individual datapoints remained above the median values for ProA and IEX, only three methods had median values greater than the median for IEX. Only one method, also based on affinity interactions, had a median HCP LRV greater than that of ProA. Therefore it seems that, in general, phase separation-based methods are not yet ready to completely replace standard chromatography operations in mAb production on a selectivity basis.

We also compared phase separation-based protein purification methods on the basis of product yield (Figure 2B). Despite significant variability as for HCP removal, almost all methods had median yields greater than 80%, with most greater than 90%. This suggests that the yield of these methods should not be a barrier to their adoption in commercial protein drug processes, where step yields of 80-90% are generally considered acceptable.

**Figure 2:**
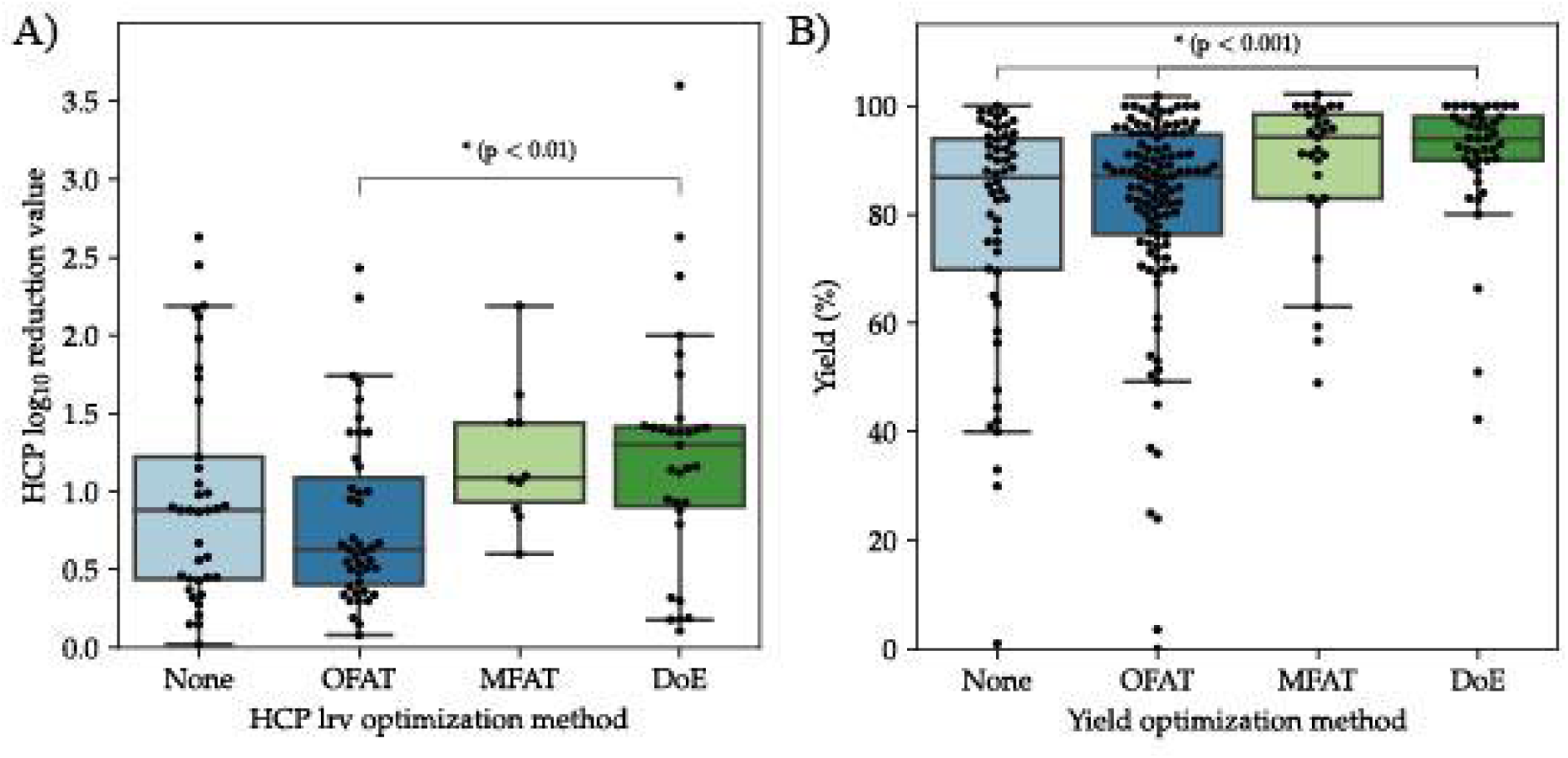
Design of Experiments results in improved method performance. A) Effect on host cell protein removal showing use of Design of Experiments significantly improves performance compared to one-factor-at-a-time optimization. B) Effect on product yield showing use of Design of Experiments significantly improves performance compared to no optimization or one-factor-at-a-time optimization. For both panels, box and whisker plots show the minimum, 25^th^ percentile, 75^th^ percentile and maximum values, respectively. Individual datapoints are shown as black circles. Following a significant one-way Kruskal-Wallis test, groups were compared by the Mann-Whitney U test with Bonferroni correction for multiple comparisons. Comparisons not shown were not significant (corrected p > 0.05).

Finally, we evaluated the methods in the dataset based on their ability to remove the other major classes of contaminants frequently encountered: high- and low-molecular weight variants of the product (HMW and LMW, respectively), endotoxin or lipopolysaccharide (LPS), and DNA. These data are summarized in Figure S11. For each of these classes of contaminants, it was found that phase separation-based methods could in some cases achieve reductions of multiple orders of magnitude, making them generally competitive with chromatography. Once again, however, more complete reporting of results is needed to facilitate evaluation of the field against current standard methods.

### 3.3. Effect of experimental design on method performance

Given the high degree of variability observed in measures of purification performance, we also investigated if the approach to experimental design was a significant factor in predicting performance in terms of HCP removal and product yield. We found that, for HCP removal (Figure 2A), the use of statistical Design of Experiments (DoE) approaches was significantly better than one-factor-at-a-time (OFAT) optimization, with a 53% increase in the mean. For yield (Figure 2B), DoE was significantly better than either no optimization or OFAT optimization, with 16.1% and 11.5% increases in the mean, respectively. Surprisingly, for both yield and HCP removal, OFAT or multiple-factors-at-a-time (MFAT) optimization did not lead to statistically significant better results than conducting no optimization.

### 3.4. Diversity of use of phase separation agents

One potential advantage of phase separation over chromatography is being able to use and independently control several selective agents with different selectivities in the same operation. Thus, to further examine the diversity in approaches to phase separation-based purification and the use of different selective modes in combination, we separated each method into its constituent selective agents and counted the number of times each agent was used either alone or in each of its possible pairwise combinations. The results of this analysis are shown in Figure 3A. We found 16 distinct agents used to effect phase separations, which were employed in 45 different combinations of either one or two agents. Thus, the pairwise combination space of agents used to purify proteins by phase separation is only approximately 1/3^rd^ explored. Furthermore, for the majority of agents, by far the most common case was that the agent was used alone (i.e., lying on the diagonal of Figure 3A). Finally, over 75% of methods using a combination of agents had 5 or fewer records. Therefore, although phase separation-based protein purification methods have a potential advantage over chromatography in their ability to independently and simultaneously select on multiple axes of protein properties, this potential remains largely unexplored or underutilized.

**Figure 3:**
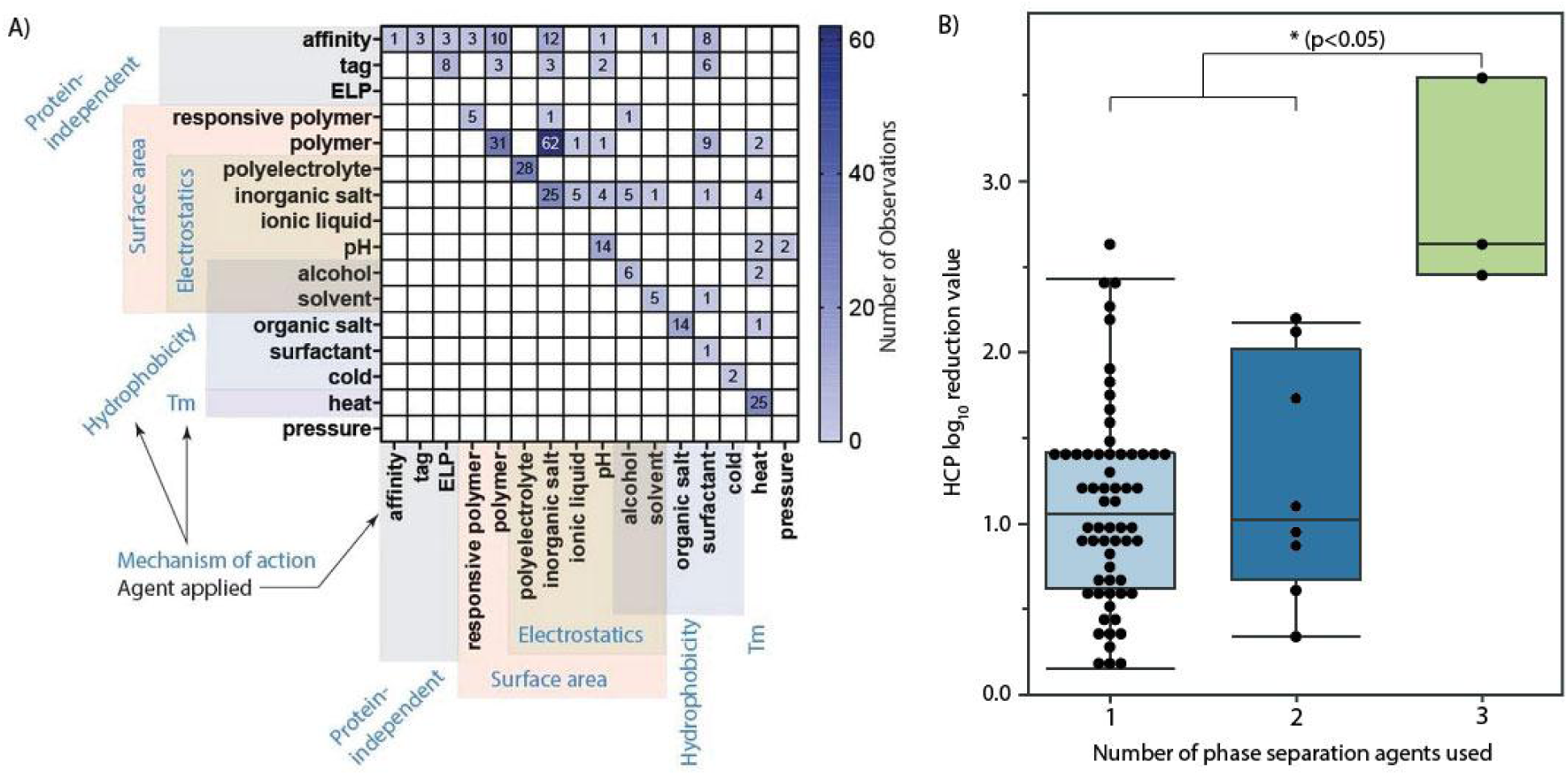
Use of selective agents in phase separation-based protein purification. (A) The pairwise combination space of selective agents used in phase separation-based protein purification is largely unexplored or underutilized. Each square shows the number of reports that utilize that agent combination. Because the matrix is symmetric, only the upper half is shown for clarity. Tag: fusions to the product protein conferring a non-native selectivity. ELP: elastin-like polypeptides either fused to the product protein or to a binding partner. Responsive polymer: polymers exhibiting sharp phase change behavior as a function of something other than polymer concentration (e.g., pH or temperature). Annotations outside the matrix show both the type of phase-forming agent used and approximate groupings based on predominant mechanisms indicated by the literature. (B) Improvement of HCP removal using an increasing number of separation agents. Box and whisker plots show the minimum, 25^th^ percentile, 75^th^ percentile and maximum values, respectively. Individual datapoints are shown as black circles. Following a significant one-way Kruskal-Wallis test, groups were compared by the Mann-Whitney U test with Bonferroni correction for multiple comparisons. Comparisons not shown were not significant (corrected p > 0.05).

Given this, we sought to determine if the use of an increasing number of separation agents in combination could, in general, lead to an increased performance in terms of host cell protein removal. We grouped phase separation methods by the number of separation agents they utilized (1 to 3) and for which HCP data was available (Figure 3B). Additionally, a method employing one or two agents was only included if at least one of those agents was also employed in other methods utilizing a greater number of separation agents (two or three agents, respectively). Although we identified only three records employing a combination of three separation agents in all the dataset, all three methods demonstrated superior performance compared to those utilizing fewer agents. This underscores the significance of delving into the combinatorial space of phase separation agents for further exploration.

### 3.5. Techno-Economic Analysis

In addition to purification performance, process economics are a key factor affecting the readiness of phase separation methods to be adopted into commercial processes. This is especially true for protein drug products, for which manufacturing costs are especially high and current adoption of phase separation methods is low. Therefore, for each record in the dataset with sufficient information, we conducted a techno-economic analysis by generating a pair of *in silico* unit operation cost models: one for the prescribed phase separation operation and another for an alternative IEX chromatography operation designed to process the same input stream. The dataset for techno-economic analysis consisted of 72 records representing 14 phase separation methods.

#### 3.5.1. Effect of process scale on cost-effectiveness

After constructing the unit operation models for each record in the dataset, we simulated each at four different scales: with phase separation or chromatography unit operations sized to yield 10, 100, 1000, or 10000 kg of purified product per year. The first three of these scales were chosen to represent reasonable current values for protein drug production because they are approximately the 25% percentile, median, and maximum of current mAb drug production scales (Figure S12). The largest scale is perhaps reasonable for a very small number of products such as insulin (Gotham et al., 2018), but was also chosen as a speculative test case that may become relevant for more products in the future (Kelley, 2007). After obtaining an estimated operating cost per kg of purified product for each model, we then obtained a measure of cost-effectiveness by further normalizing costs per kg of product by the degree of HCP removal achieved, using either the reported HCP LRV for the phase separation method or a median HCP LRV of 1.03 for IEX (see Table S2).

The first noteworthy finding of our techno-economic analysis was that, across all methods, phase separation-based protein purifications were highly unlikely to be cost-effective compared to IEX chromatography at product throughput scales of 10 kg/yr or less (Figure 4A). However, each increase in process scale showed a major increase (50% – 182%) in the fraction of phase separation models that were cost-effective compared to their chromatographic counterparts. By far the largest such increase in proportional terms occurred between the 100 and 1000 kg/yr scales (Figure 4B and C, respectively), suggesting that this range of scales may represent a rule-of-thumb “tipping point” for phase separation cost-effectiveness. However, it is also noteworthy that more than 30% of records in the dataset reported phase separation methods that were not predicted to be cost-effective even at extremely large scales of 10000 kg/yr (Figure 4D).

**Figure 4:**
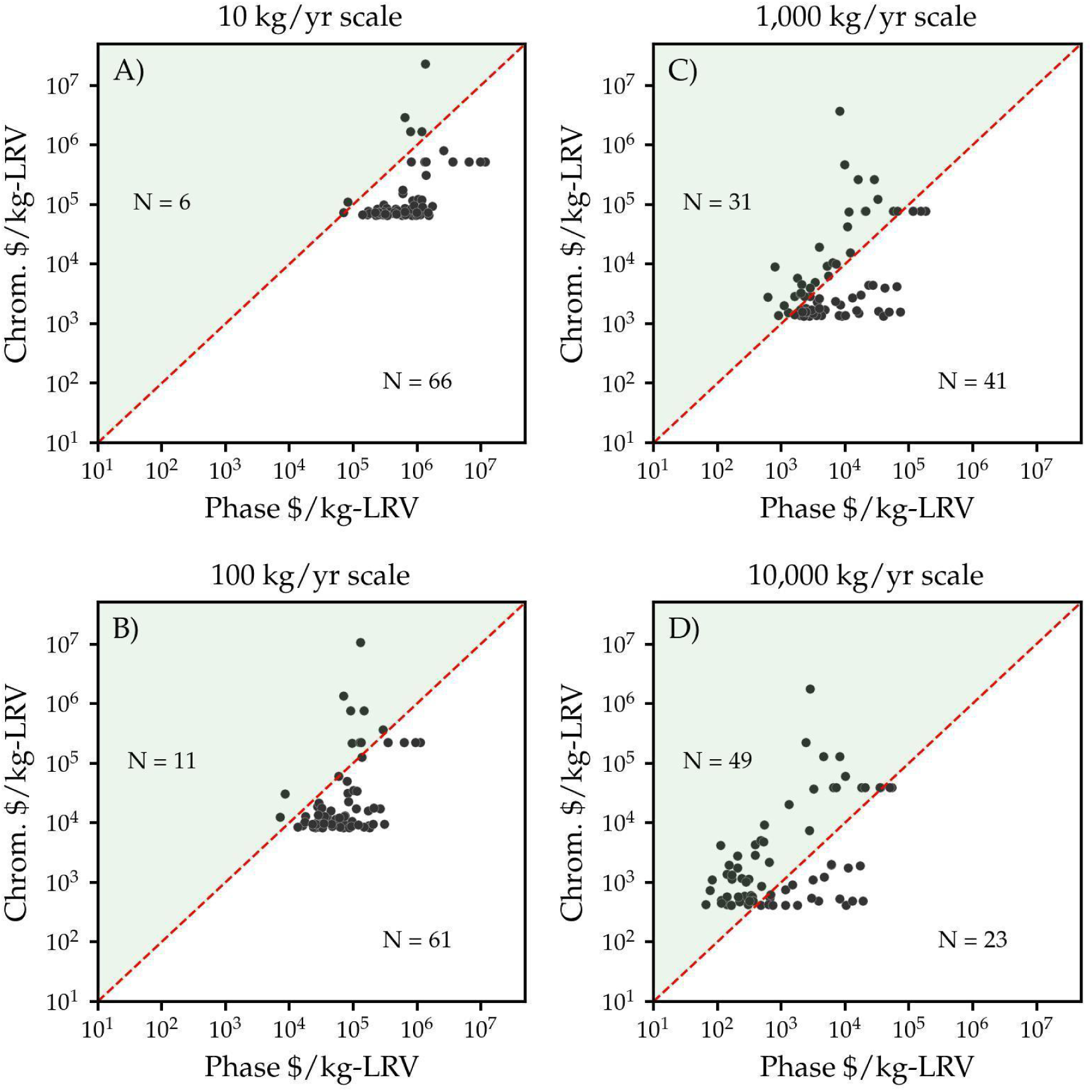
Cost effectiveness of phase separation vs chromatography. Phase separation methods become more cost-effective than ion-exchange chromatography-based alternatives as production scale increases. Panels show results at four different levels of annual product throughput. Each point represents the cost per kg of product purified normalized by the log_10_-reduction value (LRV) of host cell protein contaminants, as calculated for a given phase separation operation (x-axis) or the alternative chromatographic operation (y-axis). The red line shows equality of cost-effectiveness between chromatography and phase separation; points above the line are cost-effective compared to chromatography while those below the line are not. Annotations show the number of records in each group.

#### 3.5.2. Effect of input stream purity on cost effectiveness

Another major trend that emerged from our techno-economic analysis was that phase separation models were much more likely to be less expensive per kg of product yielded than their paired chromatographic models when the product purity in the input stream was low rather than when it was high. For example, of approximately 15 records in the dataset with an input purity of less than 1%, 100% of the phase separation models were less expensive per unit product yielded than their chromatographic alternatives at all scales above 100 kg/yr (Figure 5). By contrast, for all 72 records, the fraction of phase separation models which are less expensive per unit product than chromatography dropped to less than 80% even at the largest process scale (Figure 5B). This effect also showed a clear interaction with process scale, with initial purity showing the largest effect on cost at scales on the order of 100 kg/yr (Figure 5B, compare slopes of the curves across scales).

**Figure 5:**
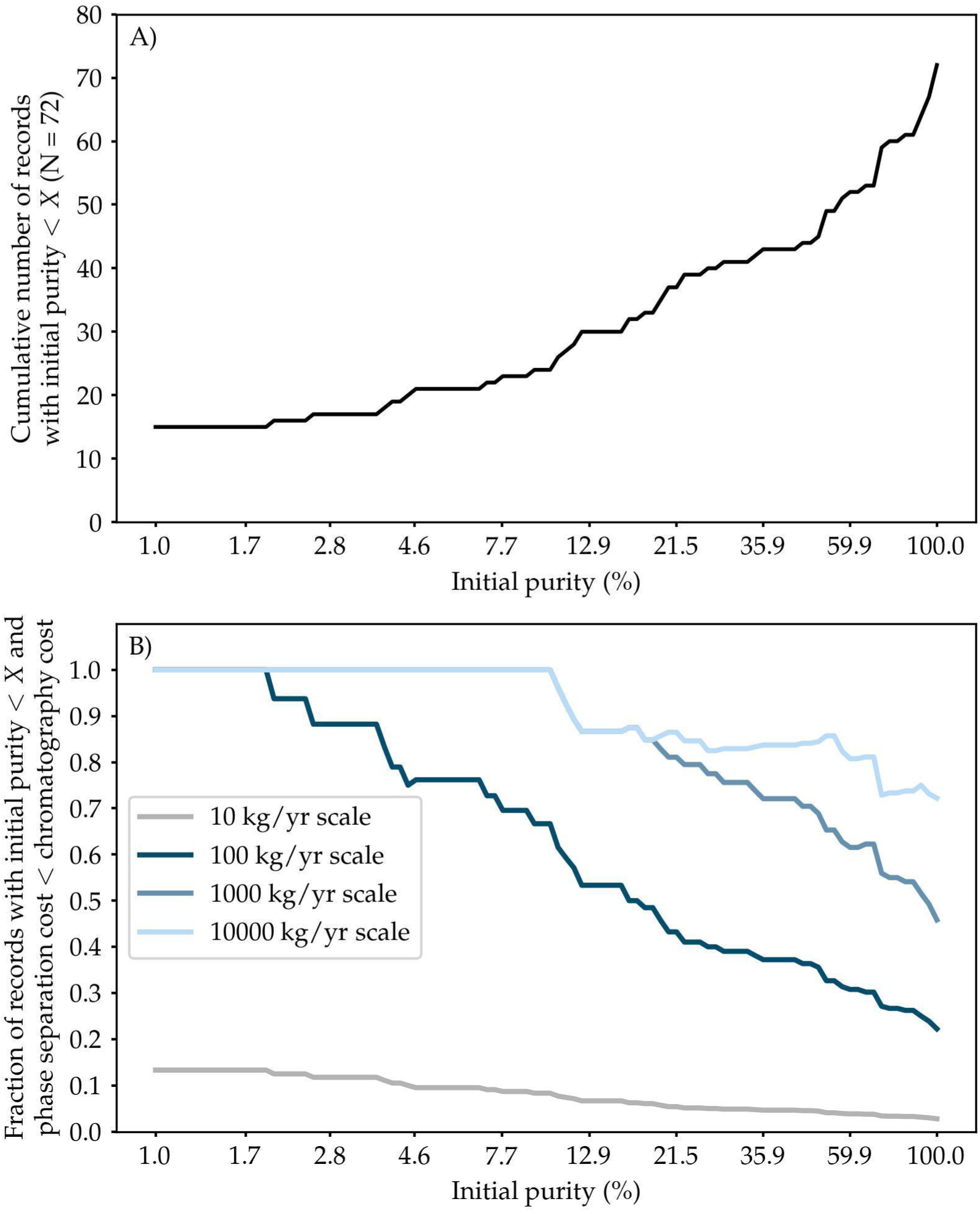
Effect of initial purity on cost-effectiveness of phase separation. A) Cumulative distribution of records in the dataset by initial purity level (i.e., purity of the product in the input stream just prior to the phase separation operation). This information is provided to assist with interpretation of the fractions in panel B. B) Cumulative distribution of records in the dataset for which total cost per kg of product purified is less for the phase separation method reported than for an alternative ion-exchange chromatography operation, as a function of initial purity and annual product throughput. As initial purity of the input stream decreases, phase separation methods become more likely to be less expensive per kg of product purified than an alternative ion-exchange chromatography operation. The fraction at each purity level considers only records with an initial purity less than that value. E.g., at 1% initial purity, the denominator is approximately 15 records (see panel A). Note that the x-axis is a logarithmic scale.

#### 3.5.3. Cost of raw materials and consumables drives process cost

Having observed two trends in the total unit cost and cost-effectiveness of phase separation purification methods vs. chromatographic alternatives, we next sought to explain the mechanisms behind these trends. We found that the best explanation for both trends across the entire dataset was a variation in the direct materials cost (from either chromatographic resin and buffers or phase-forming materials). First, we observed that as process scale increases, the cost of these direct materials makes up an increasing fraction of the total unit operation costs for both chromatography and phase separation models (Figure S13). For all chromatography models and almost all phase separation models, direct materials represent the single largest cost at the 10000 kg product/yr scale. Even at smaller scales, many of the phase separation models are dominated by these costs.

Second, we observed that there is a strong inverse relationship between initial product purity and direct materials costs per kg of product purified for chromatography, but not for phase separations (Figure 6). Moreover, direct materials costs per unit product purified decrease sharply with increasing scale for phase separation methods but much less significantly for chromatography (Figure 6, compare among panels). This is because chromatographic resins do not reap the advantages associated with economies of scales (Kelley, 2007) while bulk chemicals used in phase separations do (Qi, Wei, Roger Sathre, William R. Morrow III, Arman Shehabi, 2015). There is therefore an interaction between initial purity and process scale that affects the difference in direct materials unit costs between chromatography and phase separations. The net effect is that, as scale rises and initial purity decreases, more phase separation models fall below their counterpart chromatographic models on the direct materials unit cost axis.

**Figure 6:**
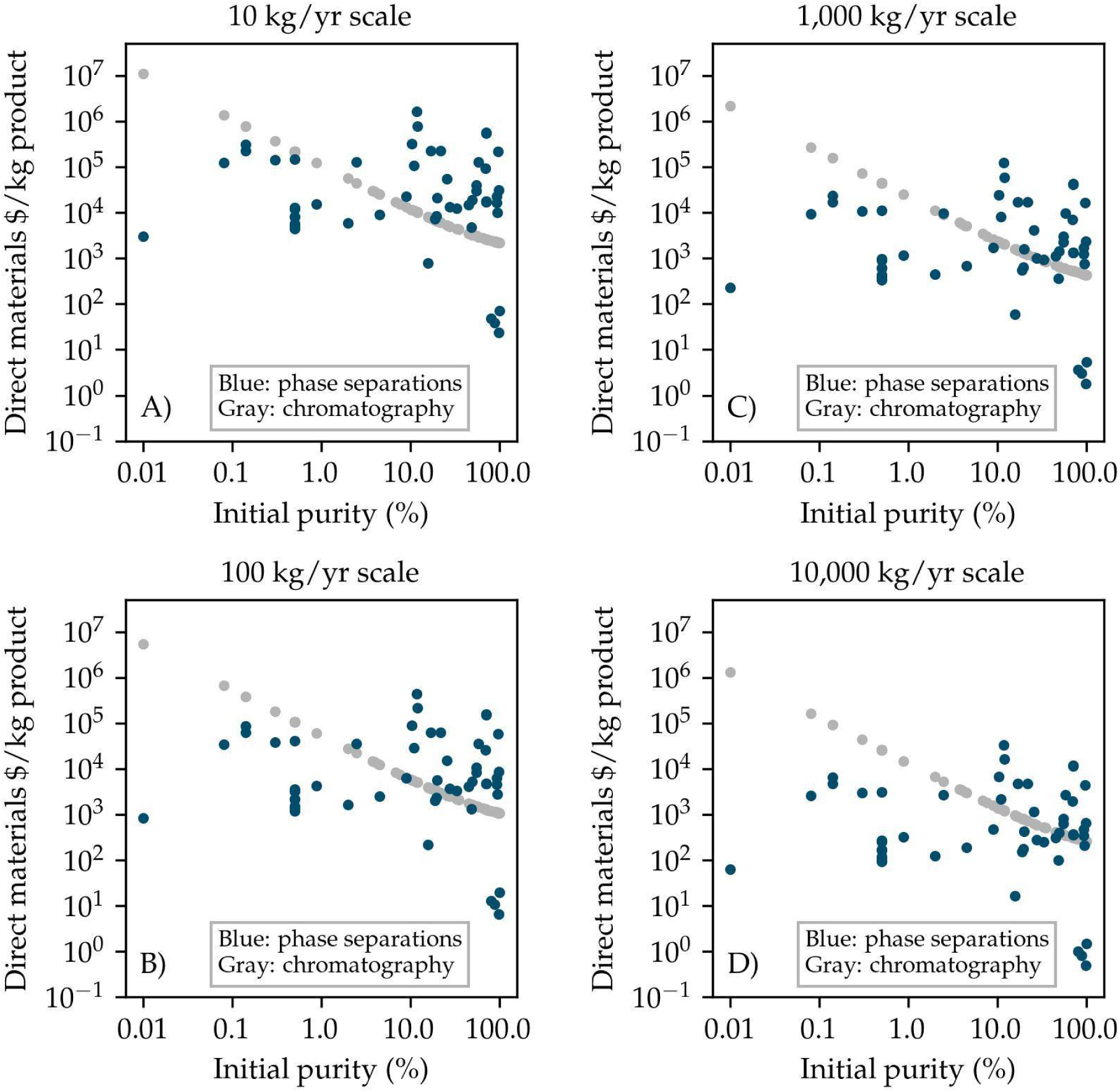
Effect of initial purity on direct material cost. Costs of direct materials per kg of product purified are inversely related to initial product purity for chromatography methods, but not for phase separation methods. Direct materials: either phase-forming materials, in the case of phase-separation methods, or resin and chromatography buffers, for an alternative ion-exchange chromatography operation. Panels show four different levels of annual product throughput. Phase separations and chromatography are shown in blue and gray, respectively.

#### 3.5.4. Direct materials usage rate as a predictor of total costs

We have established the importance of direct materials costs in explaining the trends of increasing economic favorability of phase separations over chromatography with increasing process scale and decreasing initial product purity. Next, we wanted to assess the degree to which differences in direct materials usage could also explain differences in cost among phase separation methods. To do so, we turned to the direct materials usage rate defined as the ratio of the mass of phase-forming agents and buffer components prescribed by the reaction conditions to the mass of product yielded. We found that, for all 72 records and at all four modeled process scales, total costs per kg of product purified were significantly correlated with the direct materials usage rate. Least-squares linear regressions of the log-transformed data are shown in Figure 7. Depending on scale, the direct materials usage rate alone explained between 51.4% and 58.4% of the variation in total unit operation costs per mass of product purified (R^2^ of 0.514 – 0.584). In other words, the direct materials usage rate provides a strong and simple predictor of total costs per kg of product purified across all phase separation models. It is also worth noting that the slope of the correlation increased from 0.144 to 0.32 with increasing process scale, such that at the 10000 kg/yr scale, each 10-fold increase in direct materials usage rate would produce on average a doubling (10^0.32^ ≈ 2.1) in the total process cost. Because the direct materials usage rates observed in the dataset spanned more than four orders of magnitude, this is an important effect.

**Figure 7:**
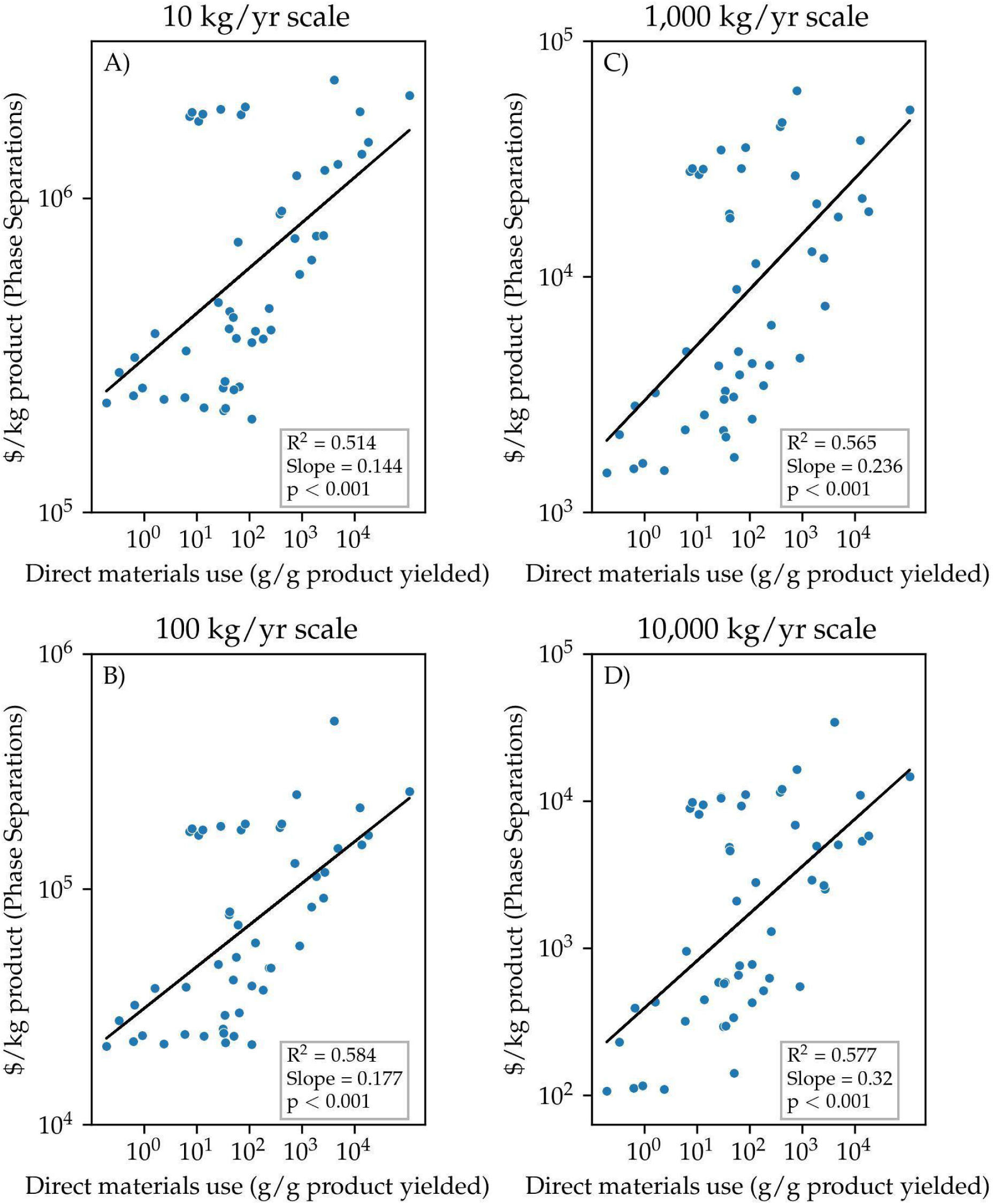
Cost of product as a function of direct material use. Cost per kg of purified product for all phase separation methods is directly related to the ratio of phase-forming materials mass to purified product mass. Black lines and inset boxes show the line of best fit and parameters for a linear regression of the log-scale data. Panels show results at four different levels of annual product throughput.

Having identified a good global predictor for the total cost per unit of product purified for all phase separation methods, we next explored the variability among and within method types for total cost and cost-effectiveness. With respect to total unit operation costs (Figure S14), we found three trends. First, many method types undergo a notable shift in cost with scale. In general, methods using no or few phase-forming materials (e.g., those based on factors such as pH and temperature) became less expensive at scale while those relying on phase-forming chemicals (e.g. inorganic salt-based precipitations) became more expensive. Second, the variability within a given method generally increased with increasing scale. Finally, the range of costs across method types was either nearly or completely overlapping at all scales. This suggests again, as for HCP removal, that there is no single best or worst phase separation method in general, but that the within-method variation is most important. With respect to cost-effectiveness ($ per kg of product per log_10_ reduction in HCPs), we note that the same themes of within-method variability applied (Figure S15), while the rank order of methods was substantially different than for cost alone (Figure S14-15).

Next, we sought to test whether direct materials usage rate could also explain the within-methods cost variations. We selected four methods that showed the highest cost variability based on Figure S14—polymer/salt ATPSs as well as precipitations based on polymers, polyelectrolytes, and inorganic salts—and used linear least-squares regression to fit the cost predictions for all examples of each method to each example’s direct materials usage rate. The results of this analysis at both the largest and smallest modeled process scales are shown in Figure S16. Of these 8 regressions, all were highly significant (p < 0.005) except that for polymer/salt ATPSs at the largest process scale, which also approached significance (p ≈ 0.15). More importantly, of the seven significant regressions, all explained more than 80% of the within-method variability in process costs, while the best explained more than 98% (R^2^ = 0.81 – 0.984). This means the direct materials usage rate almost entirely explains the within-method cost differences.

### 3.5. Implementation-Readiness of Phase Separation Methods

Finally, having compared the phase separation methods in the dataset on the basis of purification performance as well as economic performance, we sought to combine these measures to form a single assessment of the degree to which phase separation methods were currently ready to be implemented into commercial processes for protein drugs. To this end, we defined three categories of implementation readiness. The first includes methods that are not cost-effective compared to conventional IEX chromatography, as measured by cost per kg of purified product per log_10_ reduction in HCPs. These methods are therefore not generally expected to be suitable at present for adoption in protein drug processes, because adding additional separation capacity by chromatography would be more cost-effective. The second includes measures that are cost-effective compared to IEX chromatography, and therefore may be considered as worthwhile supplements or partial replacements to an existing chromatography train. Finally, the third category contains methods that have the potential to entirely replace at least one standard chromatographic unit operation in a downstream process, because they are both cost-effective compared to IEX and have HCP LRVs greater than the median for IEX.

First, we compared implementation readiness as a function of the year of publication of each study in the techno-economic analysis dataset (N = 72) as well as of process scale (Figure S17). As an encouraging sign for the field, we observed that the rate of “implementation-ready” phase separations has been increasing over time. For example, at the 10 kg/yr scale, only one such example was published in the 16 years between January 2000 and December 2015, while 5 were published in the 6 years between January 2016 and January 2022.

Next, we sought to determine whether differences in the method used to induce phase separation were associated with differences in implementation readiness (Figure 8). We found that a strong association did indeed exist. At the 10 kg/yr and 100 kg/yr production scales, respectively, only 3 and 5 of the 14 method types had any implementation-ready examples. While most methods had at least one implementation-ready example at the 1000 kg/yr scale, we note that the great majority of current protein drug processes operate below this scale (Figure S12). Notably, the methods most likely to be cost-effective or potential chromatography replacements were all precipitations, and included those based on polyelectrolytes, inorganic salts, polymers, and the combination of heat, inorganic salts, and extreme pH. For more details on which methods are implementation-ready or not, refer to Table S6.

**Figure 8:**
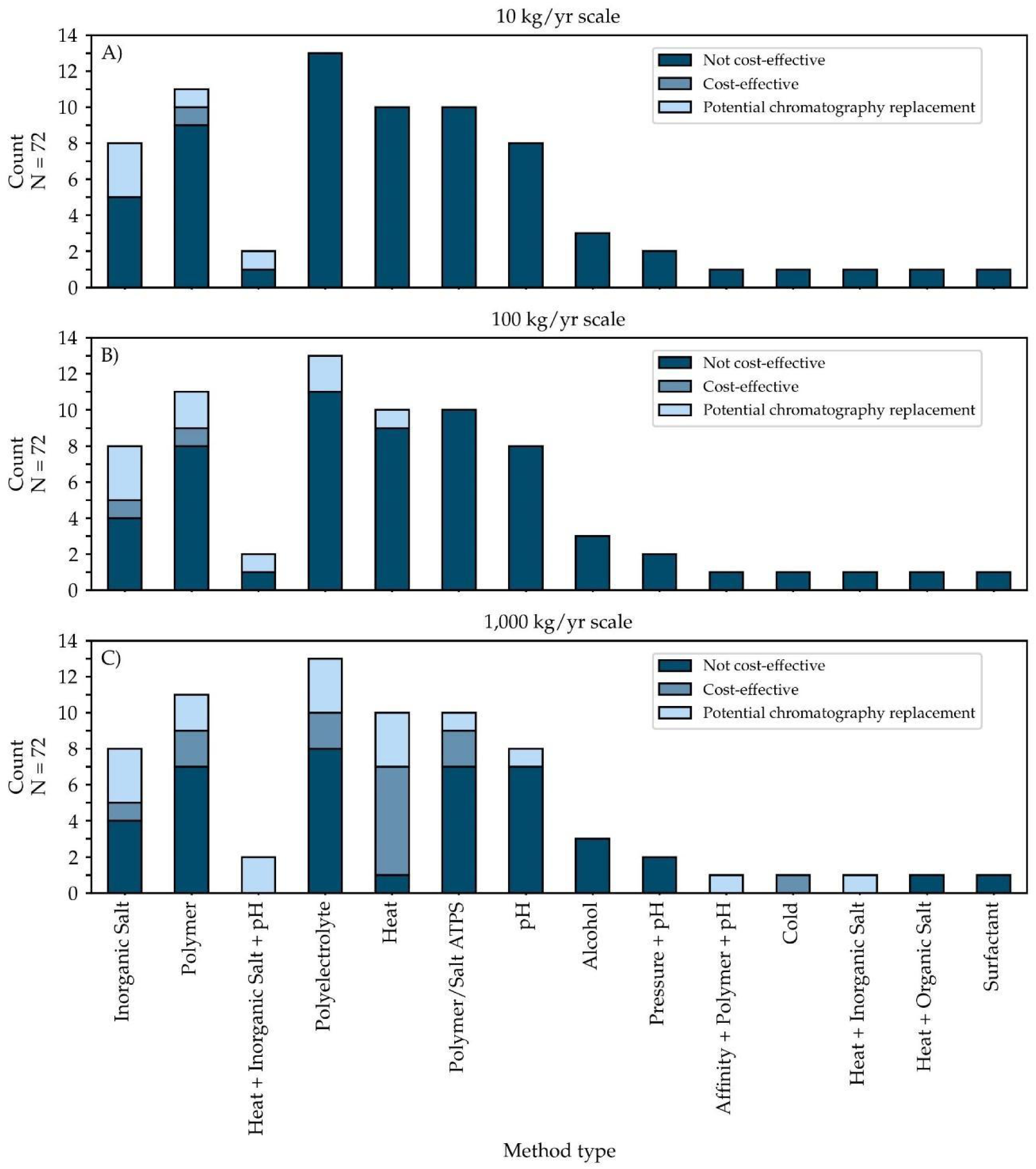
Implementation readiness as a function of method type and annual product throughput. Methods are precipitations unless ending in ATPS (aqueous two-phase separation). For further information on the definitions of technology readiness categories, refer to the text.

Then, because the approach to experimental design to optimize the reaction conditions was found to significantly affect purification performance in terms of product yield and HCP removal, we tested whether the same factors were associated with increased likelihood of implementation-readiness. In general, it did not appear that more advanced optimization methods (e.g., MFAT and DoE approaches) were associated with greater rates of technology readiness than simple OFAT optimization when optimizing either for HCP removal (Figure S18) or for product yield (Figure S19). Furthermore, optimization of these variables by any method was not clearly better than not optimizing them at all, except at large scales (e.g., 1000 kg/yr, Figure S18C and S19C). We note that there were no examples in the entire analysis dataset (N = 290) where the authors optimized directly for cost or for lowering the direct materials usage rate here shown to drive cost.

## 4. Discussion

Methods of protein purification based on modulating solubility among different chemical phases (i.e., phase separations), when used in place of standard alternatives such as chromatography, have the potential to simplify protein manufacturing and reduce its costs. This is especially important in the area of protein drug manufacturing, where process costs and complexity are highest. However, while phase separations are in common use in other areas such as industrial enzyme production, they have not been adopted in protein drug manufacturing outside of a small number of cases (Dos Santos et al., 2017; Thömmes and Etzel, 2007). Our purpose in this study was to identify the reasons for this lack of uptake, to assess the current readiness of technologies in the field for implementation in protein drug processes, and to offer evidence-based guidelines for future work in the field to help increase implementation-readiness.

### 4.1. Operating Cost is a Major Barrier to Implementation

Previous reviews in the field have primarily offered two reasons for the lack of uptake of phase separation methods in protein drug manufacturing: lack of mechanistic and process engineering knowledge (Dos Santos et al., 2017; Gagnon, 2012; Hekmat, 2015; Martinez et al., 2019; Soares et al., 2015); and insufficient or inconsistent selectivity (Martinez et al., 2019). Meanwhile, the economic aspects of using phase separation vs. standard alternatives such as chromatography in the protein drug context are usually treated only superficially. In fact, it is often taken almost axiomatically that phase separation methods will be less expensive than chromatography. Although some recent studies do treat this question at some depth, they only consider processing scales much larger than those in common use for protein drugs (Rosa et al., 2011) and/or only one case study of a particular phase separation method and product protein (Decker et al., 2020; Rosa et al., 2011).

In contrast to previous work in the field, we present the first in-depth techno-economic analysis of the full range of demonstrated use-cases for phase separations in protein purification. To our surprise, and in contrast with conventional understanding in the field, we found that phase separation methods are in fact frequently economically unfavorable vs. standard alternatives under a wide range of process conditions applicable to most of the currently marketed protein drugs. Furthermore, although our definition of cost-effectiveness incorporates the degree of HCP removal achieved, this unfavorable economic comparison of many phase separation methods to chromatography is not due solely to a lack of selectivity, because it persists when accounting only for total costs rather than cost-effectiveness (Figure 5B).

In general, we found that the most favorable process conditions for phase separation methods to be cost-effective compared to chromatography were large scale (Figure 4), especially greater than 1000 kg/yr, and low initial purity of the product in the input stream (Figure 5B). Moreover, these two factors have an interaction, where initial purity is most important at moderately large scales (e.g., 100 and 1000 kg/yr) (see Figure 5B).

### 4.2. The direct materials usage rate can predict process cost

After finding that both initial product purity and process scale strongly influence the cost of phase separation methods compared to chromatography, we sought mechanistic explanations for these effects. We found that the common factor explaining both trends was changes in costs of direct materials (i.e., phase forming materials or chromatography resins and buffers) per unit of product purified. This is established by three facts. First, as process scale increases, these direct materials costs come to dominate total costs for both phase separations and chromatography (Figure S13). This is a well-known effect in downstream processing, as these materials scale more closely with the amount of product processed than do other costs such as equipment or labor (Farid, 2017). Second, as process scale increases, the costs of direct materials per unit of product purified falls sharply for phase separations but much less strongly for chromatography (Figure 6). This is due to the fact that chromatographic resins do not enjoy significant economies of scale in these scale regimes (Kelley, 2007), whereas raw materials used in phase separations more commonly do (Farid, 2017; Qi, Wei, Roger Sathre, William R. Morrow III, Arman Shehabi, 2015). Third, there is a strong dependence of the direct materials costs for chromatography, but not for phase separation, on the purity of the input stream (Figure 6). This is because chromatography involves a direct interaction between the resin and both the product as well as at least some contaminants, so that the resin mass required is directly related to the mass of protein in the input stream and inversely related to its purity. In contrast, many materials used to induce phase separations are based on changing the properties of the solvent rather than on interacting with proteins directly (Arakawa and Timasheff, 1985). Therefore, the required mass of these materials scales directly with the process volume and inversely with initial product concentration, but not necessarily with the protein mass or purity of the input stream. While this difference in scaling behaviors has been noted before (Dos Santos et al., 2017; Oelmeier et al., 2013), neither its implications for initial product purity nor its detailed impact on total process costs have previously been reported.

We noted above that cost-competitiveness with chromatography can be an important barrier for implementation of phase separation methods and that direct materials costs explain much of the differences in cost between phase separations and chromatography. However, it is not reasonable that every new phase separation study should contain a detailed economic analysis to determine its cost-effectiveness in these ways. Therefore, we sought a simple factor that could predict process costs both among different types of phase separation methods and among different examples of the same type of method. We found this factor in the direct materials usage rate. This simple measure, which does not even consider variations in the costs of different types of phase-forming materials, quantitatively explains cost variations both across phase separation method types and within a given method type. Therefore, in lieu of conducting a complete economic assessment, future work in phase separation method development should at least consider the direct materials usage rate as an easily-calculated proxy for economic viability. Of course, this metric does not necessarily apply to methods that use strictly or primarily non-chemical means such as temperature or pressure to induce phase changes.

### 4.4. Some phase separation methods have Demonstrated Implementation-Readiness for protein drug processing

Considering both cost-effectiveness (a combined measure of process cost, yield, and contaminant removal) and total HCP removal capabilities, we classified reported phase separation examples into three categories of implementation-readiness. Broadly, the categories correspond to examples of phase separations that are not worth implementing in a protein drug process, are worth implementing as a supplement to chromatography, or are worth implementing as a replacement for chromatography. Interestingly, we found that at the 10 kg/yr and 100 kg/yr process scales, only a few of the many different types of phase separation methods were ready to supplement or replace chromatography. At these small to moderate scales, the methods most ready for implementation were precipitation employing polyelectrolytes, inorganic salts, polymers, and the combination of heat, inorganic salts, and extreme pH. At the 1000 kg/yr scale, the success rate for polymer/salt ATPSs also rose notably.

## 5. Conclusions

We can summarize our findings as a set of guidelines for future work in the phase separations field. Researchers should not take for granted that phase separation methods will be less expensive than chromatography. And while one may use the direct material usage rate as a predictor of economic viability (Figure 7), a full techno-economic analysis of new methods compared to chromatography is needed to make precise conclusions. Also, when developing new phase separation methods, researchers should take advantage of modern optimization techniques such as Design of Experiments (DoE) because their use significantly improves method performance. Likewise, a new method should minimize the direct materials usage rate since it has similar/more impact than HCP removal on method implementation readiness.

Notably, implementation readiness may be easier to achieve if using a combination of phase-forming agents since they result in improved method performance (Figure 3b) and lower direct material usage rates (when leveraging methods with low material usage such as pH or heat precipitations). As such, new methods should explore the combination of multiple phase-forming agents with different selective modes. Finally, near-term efforts to implement phase separation methods in protein drug processes should focus primarily on cases with large process throughputs (e.g., 100s of kg/yr or more) where a phase separation can replace or supplement a chromatography step that receives relatively low-purity input material.

## Author Contributions

JSD and MDL conceived the study. JSD and UY reviewed and analyzed publications. JSD conducted analyses. JSD, UY, RMM and MDL prepared and reviewed the manuscript.

## Acknowledgements

We gratefully acknowledge funding from NIH grant 3R61AI140485. U. Yano was in part supported by the Takenaka Scholarship Foundation. We would also like to acknowledge N. Cheatwood for his help in compiling manuscripts.

