## Supplemental Materials for "Precipitation and Extraction Methods for Protein Purification: A Meta-Analysis of Purification Performance and Cost-Effectiveness"

### 1. Supplemental Methods

#### 1.1. Literature Search

The term used to search the PubMed database was:

“((((("precipitat*"[Title/Abstract] OR ("aqueous"[Title/Abstract] AND ("extract*"[Title/Abstract] OR "phase"[Title/Abstract]))) AND ("yield*"[Title/Abstract] OR "recover*"[Title/Abstract]) AND ("remov*"[Title/Abstract] OR "clear*"[Title/Abstract] OR "reduc*"[Title/Abstract] OR "purity"[Title/Abstract] OR "separat*"[Title/Abstract]) AND ("bioprocess*"[Title/Abstract] OR "purifi*"[Title/Abstract] OR "biomanufactur*"[Title/Abstract]) AND ("antibod*"[Title/Abstract] OR "mAb"[Title/Abstract] OR "protein*"[Title/Abstract] OR "enzyme*"[Title/Abstract])) NOT "Preparative biochemistry biotechnology"[Journal]) AND 2000/01/01:3000/12/31[Date - Publication] AND "English"[Language]) NOT "Review"[Publication Type]) AND (english[Filter])”

#### 1.2. Reviewed Features

**Table S1: Definitions of features extracted from the dataset.**

| Feature | Definition |
|---|---|
| Product name | The common name (e.g., “penicillin acylase”), gene name (e.g., “pac”), or a generic name (e.g., “mAb”) for the product protein. |
| Product gene source | The common name (e.g., “cow”) or scientific name (e.g., “Bos taurus”) of the organism from which the gene encoding the product protein was derived. |
| Industrial context | The major area of use for the product or purification process. E.g., medical products, industrial agents, etc. |
| Host | The common name (e.g., “CHO”) or scientific name (e.g., “Cricetulus griseus”) of the organism from which the product was purified. |
| Method type | A categorization of the phase separation unit operation as either precipitation or liquid-liquid extraction, followed by the type of separating agents used. E.g., precipitation: heat or LLE: polymer/salt |

| | |
|---|---|
| | ATPS. Some records contain a combination of two or more basic method types, separated by commas. |
| Method description | A quotation from the publication describing the general methods used to achieve the phase separation operation. |
| Recipe | A brief description detailing, as far as possible, the chemical species, mass composition, temperature, pH, and time relevant to the phase separation unit operation. E.g., “5 mg protein, 5 g total ATPS, 2 mM Triton X-114-Cu(II), 0.5 mM HM-EOPO polymer, 10 C, 5 hr” |
| Process stage | A general categorization of where the phase separation unit operation was used in the context of a complete purification process. E.g., “capture” for operations used on crude lysates or “polishing” for operations used on nearly-pure proteins. |
| Sequence | The complete sequence of unit operations reported in the study, beginning with the product harvest step. The phase separation unit operation relevant to a given record is indicated by an “X”. If other phase separation operations are used in the same sequence and detailed in separate records, they are indicated by numerals corresponding to the numbering of the records in the publication. |
| Yield | The recovery of product from the phase separation unit operation, expressed as a percentage. When possible, mass-based measures were used. When only measures based on biological activity were reported, they were used unless it was clear that the ratio of activity to product mass was substantially affected by the operation. |
| Initial purity | The purity of the product before the phase separation unit operation, on a mass per mass basis. When possible, the measures used were relative only to protein-based contaminants. |
| Final purity | The purity of the product after the phase separation unit operation, on a mass per mass basis. When possible, the measures used were relative only to protein-based contaminants. |
| Purification factor | The ratio of final purity to initial purity. Or, if mass-based purity measures were not reported, the ratio of final specific activity to initial specific activity. |
| HCP LRV | The $\log_{10}$ reduction value in host cell protein contaminant mass achieved by the phase separation unit operation. E.g., $\log_{10}(HCP_{initial} / HCP_{final})$. |
| DNA LRV | The $\log_{10}$ reduction value in DNA contaminant mass achieved by the phase separation unit operation. |

| | |
|---|---|
| LPS LRV | The $log_{10}$ reduction value in LPS (i.e., endotoxin) contaminant mass achieved by the phase separation unit operation. |
| HMW LRV | The $log_{10}$ reduction value in HMW (i.e., high molecular weight, or aggregate) contaminant mass achieved by the phase separation unit operation. |
| Concentration factor | The ratio of product mass per volume before vs. after the phase separation unit operation, taking into account any necessary auxiliary operations such as resuspension of a precipitate. |
| Recycling | The percentage of a given reagent used in a phase separation operation that was reported to be recovered and reused in subsequent cycles. |
| Optimization method | The type of experimental design used to optimize the conditions of the phase separation unit operation. Takes values of: None reported; OFAT (One Factor at a Time experimentation), i.e., when each factor is optimized independently; MFAT (Multiple Factors at a Time experimentation), i.e., when multiple factors are varied and optimized simultaneously; or DoE (Design of Experiments), i.e., when MFAT experimentation was supplemented by the use of statistical principles to select the set of experiments performed. |
| Theoretical model | Whether any kind of theoretical model or simulation study was directly used in selecting the conditions of the phase separation unit operation. Takes values of “yes” or “no”. |
| Reagent phase data | Whether data are reported or referenced that describe the partitioning of the reagents used in the phase separation unit operation, e.g., a binodal curve for ATPS. Takes values of “yes” or “no”. |

#### 1.3. Equations and analyses applied in standardizing data from publications

$LRV = log_{10}(fold\ change) = log_{10}\left(\frac{100}{(100-Percentage\ Decrease)}\right)$. Additionally, HCP fold changes were calculated from initial and final purities and yield as: fold change = $\frac{\left(\frac{(1-Purity_{initial})}{(Purity_{initial})}\right)}{\frac{(Yield-(Purity_{final}\times Yield))}{Purity_{final}}}$.

When relevant quantitative data were shown in figures but not reported in the text, figures were digitized using WebPlotDigitizer to extract numerical data (Rohatgi, 2021). Figures that were not 2D graphs (e.g., SDS-PAGE gels, surface plots) were not analyzed.

All quantitative measures relating to the reduction of a contaminant were reported in absolute terms. In some cases, this required adjusting reported values based on the product yield. For example, if HCP levels before and after a phase separation operation were given in parts per million (ppm), then $HCP\ LRV\ =\ log_{10}\left(\frac{ppm_{initial}\times 100}{ppm_{final}\times \%\ Yield}\right)$.

For concentration factors, in the case of ATPS operations for which no mass-based concentration factor could be determined, an approximate concentration factor was calculated based on the product yield, the volume ratio of the phases, and the fraction of initial system volume made up by the product-containing component. In this case, the simplifying assumption was made that all components had the same density. For example, if an ATPS system initially contained 20% w/w product-containing lysate, the product separated to the top phase with a 90% yield, and the phase ratio was 1:1, the approximate concentration factor would be $\frac{\frac{\frac{90}{100}}{\frac{1}{2}}}{\frac{100}{20}}$.

### 1.4. Techno-economic Analysis

For phase separation operations, parameters determined from each record included the yield of product relative to the input amount, the initial product mass concentration, the host cell protein $\log_{10}$ reduction value, the concentrations of each reagent on a mass per volume or mass per mass of product basis, and the reaction duration and temperature. Materials required to titrate the pH of solutions were neglected. When initial product concentration could not be determined, an initial total protein concentration of 5 g/L was assumed and the initial product concentration was calculated from the known initial purity. When reaction temperatures and durations were not reported, they were assumed to be 25 °C and 1 hour, respectively. Base prices for reagents were obtained from commercial supplier quotes at the largest scale available (generally 2 to 25+ kg), using the highest grade of reagent available (generally pharmacopeial grade).

For chromatography, input parameters from each record were the initial product purity and the initial total protein concentration (i.e., initial product concentration divided by initial purity fraction, with initial product concentrations determined or assumed as for phase separation operations). Product yield was assumed to be 95%. Resin volumes were sized to accommodate initial binding of 100% of product and 50% of contaminants. All chromatography operations were modeled in bind-and-elute mode.

For both chromatography and phase separations, we considered only operating expenses for the relevant unit operation; capital costs and operating costs not directly related to the unit operation were not considered. Both cases included three major cost categories. First we considered facility costs, i.e., maintenance, insurance, taxes, and overhead costs resulting from both primary equipment and auxiliary equipment necessary to complete the operation. Primary equipment was assumed to be stainless steel blending tanks for phase separation operations or stainless steel columns for chromatography. Second, we considered labor costs. Labor was priced at $80.50 per hour total cost and it was assumed that 1 hour of labor was required per hour of unit operation duration per parallel equipment train. Additionally, 0.25 hours of QC were assumed per hour of labor, at the same hourly rate. The third cost category was for materials. Materials costs included resins and buffers for chromatography, phase-forming reagents and buffer components for phase separation methods, and cleaning agents for both. Economies of scale in materials prices were accounted for using a standard exponential scaling equation with an exponent of -0.56 (Qi, Wei, Roger Sathre, William R. Morrow III, Arman Shehabi, 2015) to determine a large-scale unit price based on the unit price and mass obtained from supplier quotes and the mass of each material required in the model per year. Water for injection (WFI), cleaning agents, and buffers were adjusted in the same way except that base prices and scales were sourced from the literature (Farid, 2017). Resin costs were not subjected to economies of scale, in line with previous analyses of large-scale protein drug production (Kelley, 2007). Finally, for phase separation operations only, utilities costs were included to account for electrical heating or cooling to any temperatures prescribed by the method description other than 25 °C. Chromatography was assumed to occur at 25 °C.

Additional parameters relevant to US-based pharmaceutical biotechnology were obtained from SuperPro Designer (Intelligen, Scotch Plains, NJ). These included size-based prices for primary equipment, maximum equipment sizes, and coefficients relating primary equipment costs to auxiliary equipment costs and facility-based operating costs.

### 2. Supplemental Results

#### 2.1. Classification of the dataset

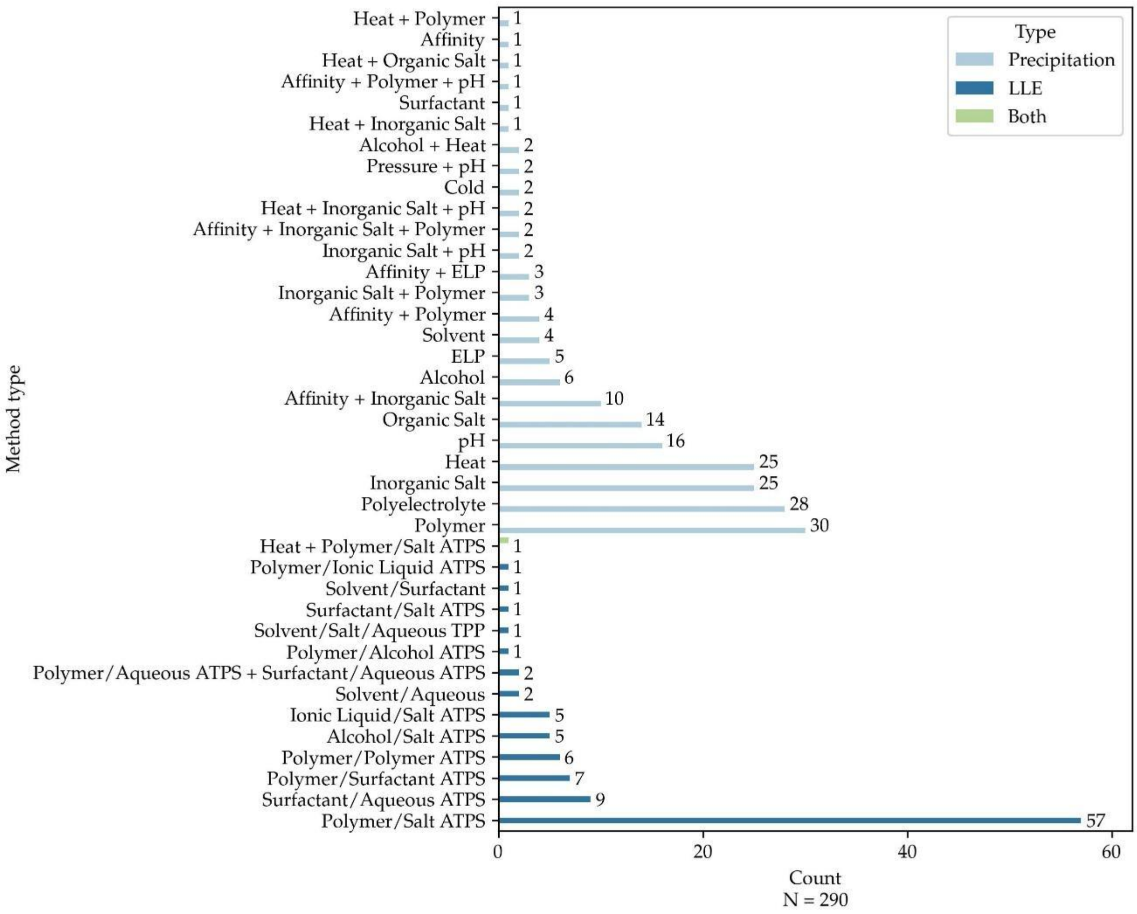


**Figure S1: Number of reports per type of phase separation method.** Many different phase separation-based protein purification methods are in use, including many using more than one kind of phase-forming agent, but most have low numbers of reports. LLE: liquid-liquid extraction. ATPS: aqueous two-phase separation. TPP: three phase partitioning. ELP: elastin-like polypeptide. Bar annotations show the number of records in each group (N=290).

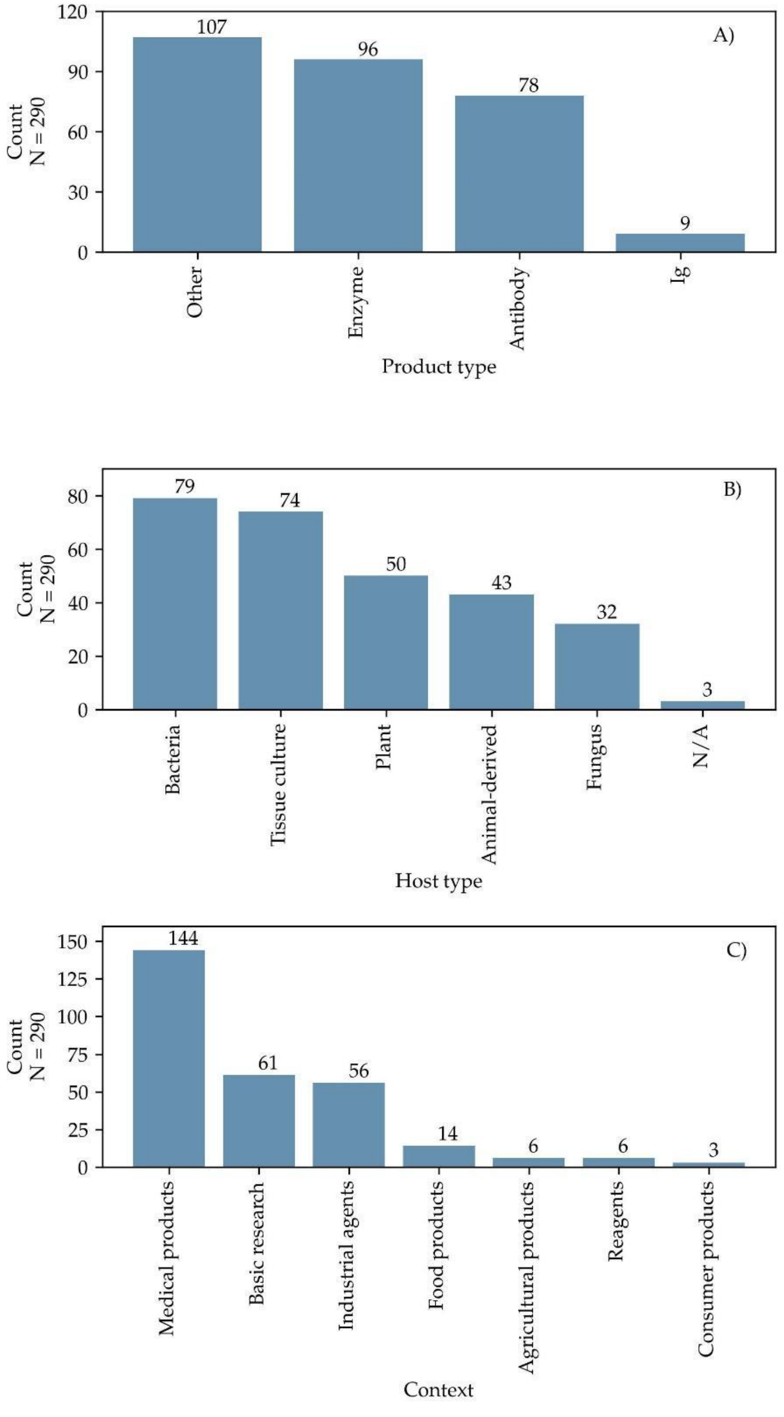


**Figure S2: Dataset classification by product type, host organisms, and research application.** A) Broad categories of protein products represented in the study. B) Host organisms, i.e., the source of the background proteins from which the product was purified. N/A: not applicable, e.g., because pure proteins were studied. C) Research contexts, i.e., the main area of purported application for the product or process.

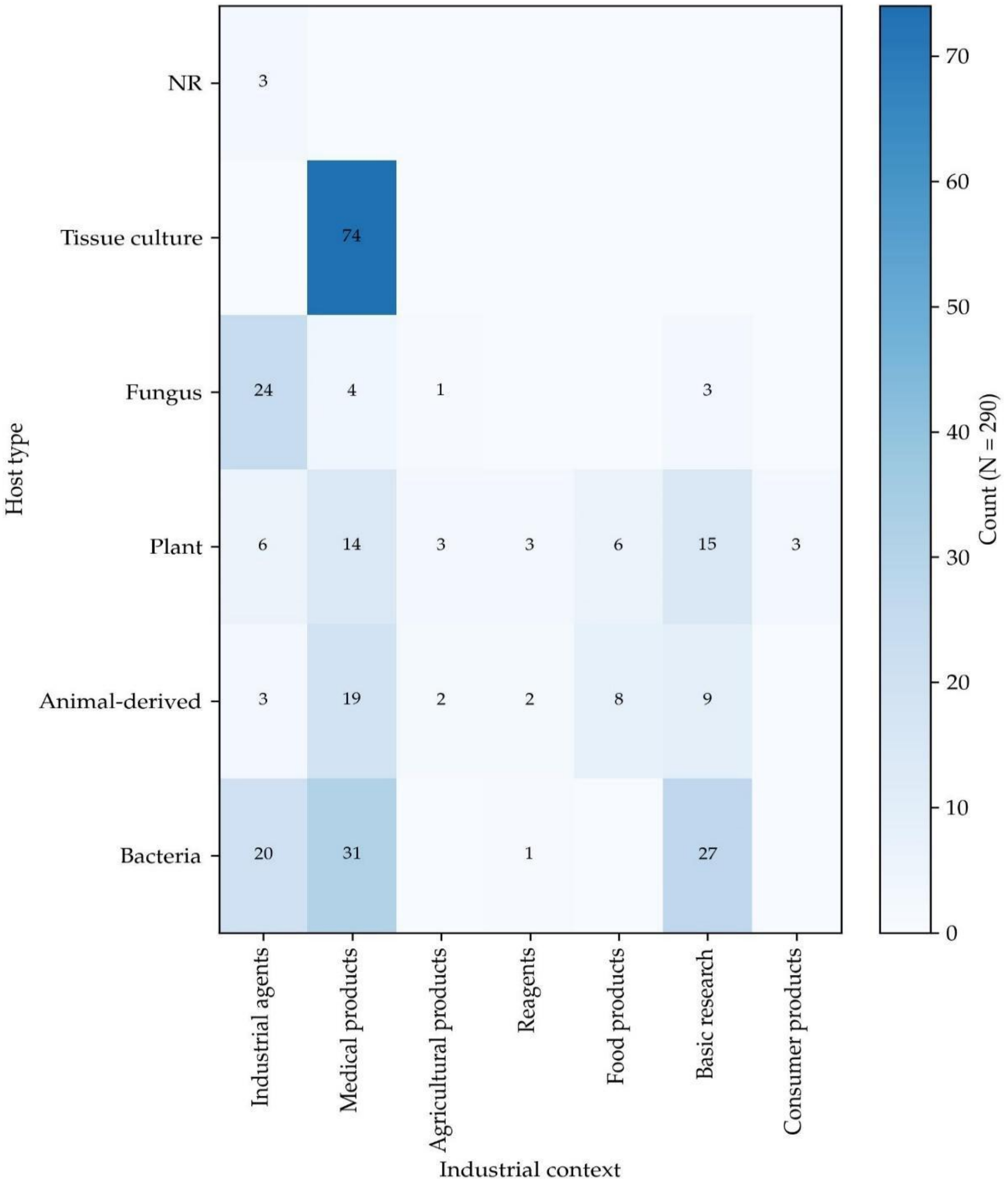


**Figure S3: Dataset representation by combination of host and research application.** NR = not reported. For further information on categorization of research contexts, refer to Chapter 4.

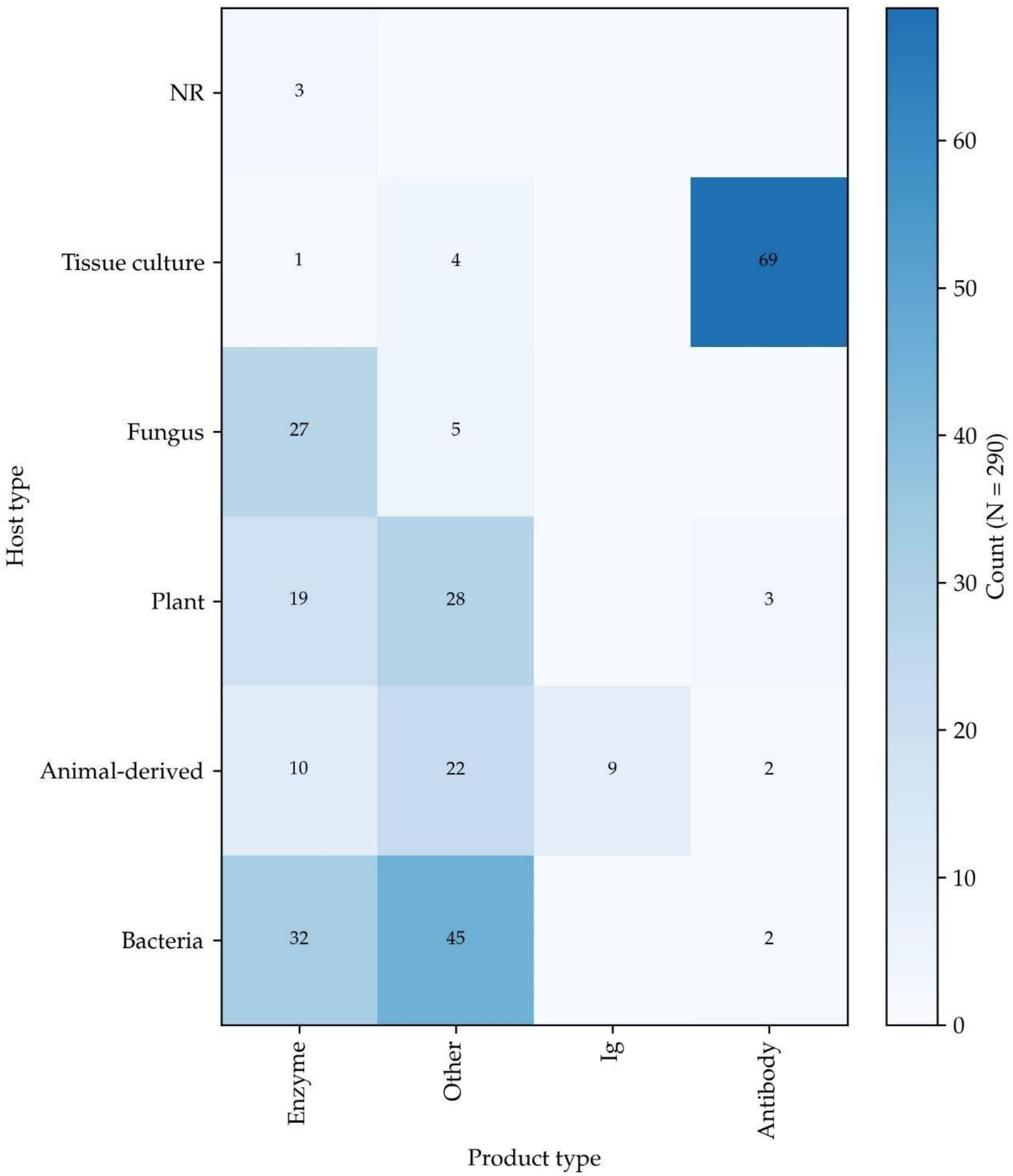


**Figure S4: Dataset representation by combination of host and product type.** NR = not reported. For further information on product categorization, refer to Chapter 4.

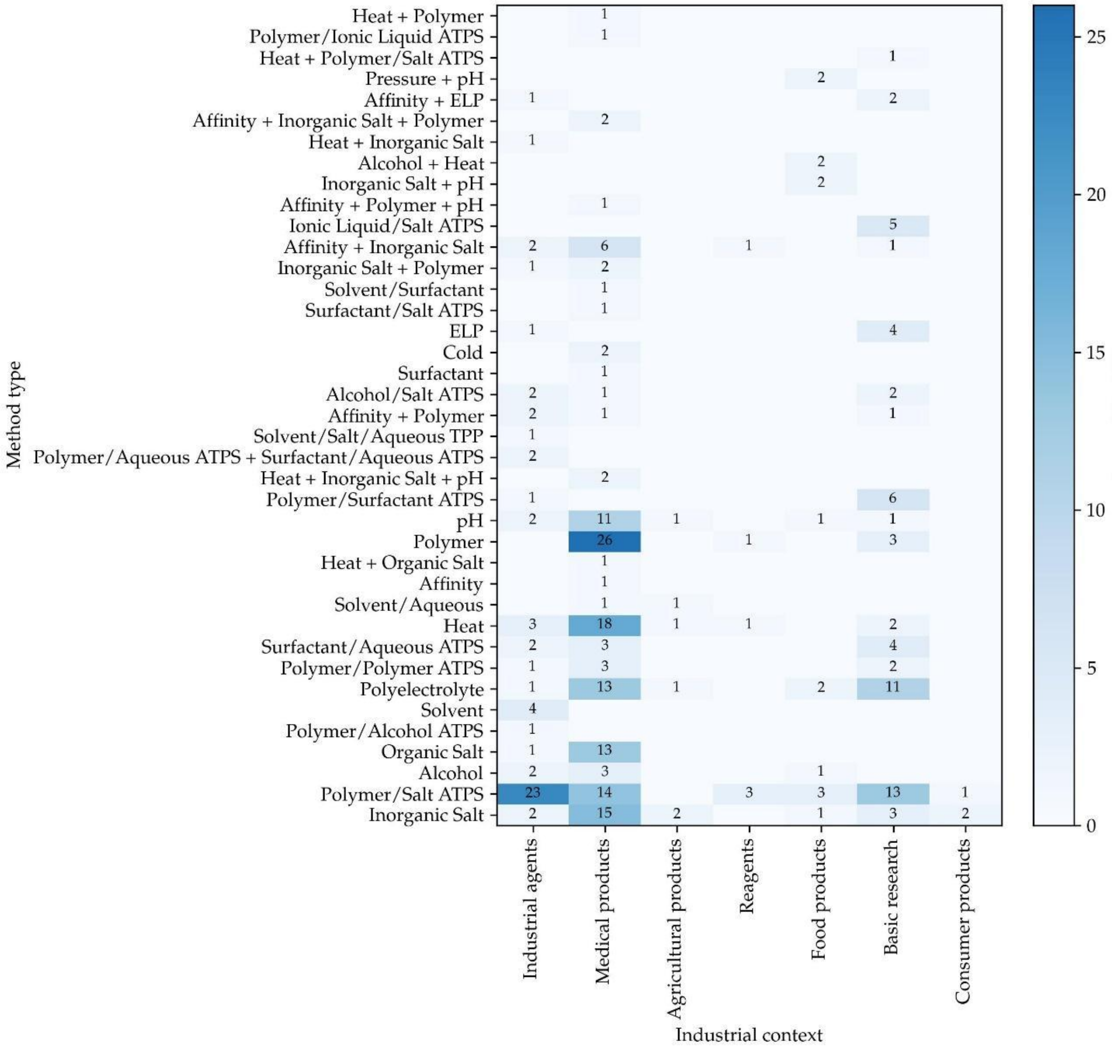


**Figure S5: Usage of different phase separation methods as a function of research application**. Method types are precipitations unless ending with ATPS (aqueous two-phase separation) or TPP (three-phase partitioning). For further information on categorization of research context, refer to Chapter 4.

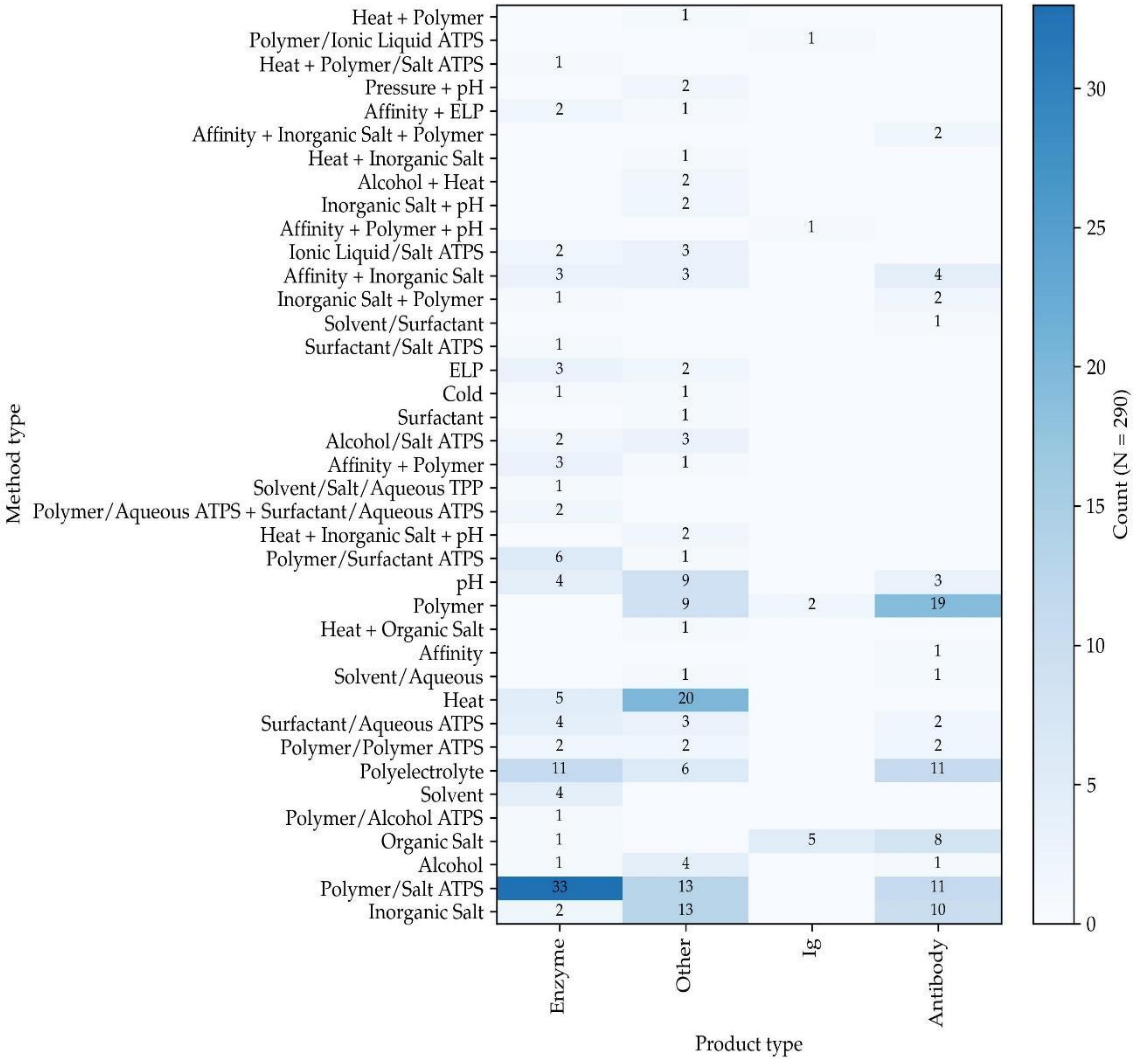


**Figure S6: Usage of different phase separation methods as a function of product type.** Ig = Immunoglobulin. Method types are precipitations unless ending with ATPS (aqueous two-phase separation) or TPP (three-phase partitioning). For further information on product categorization, refer to Chapter 4.

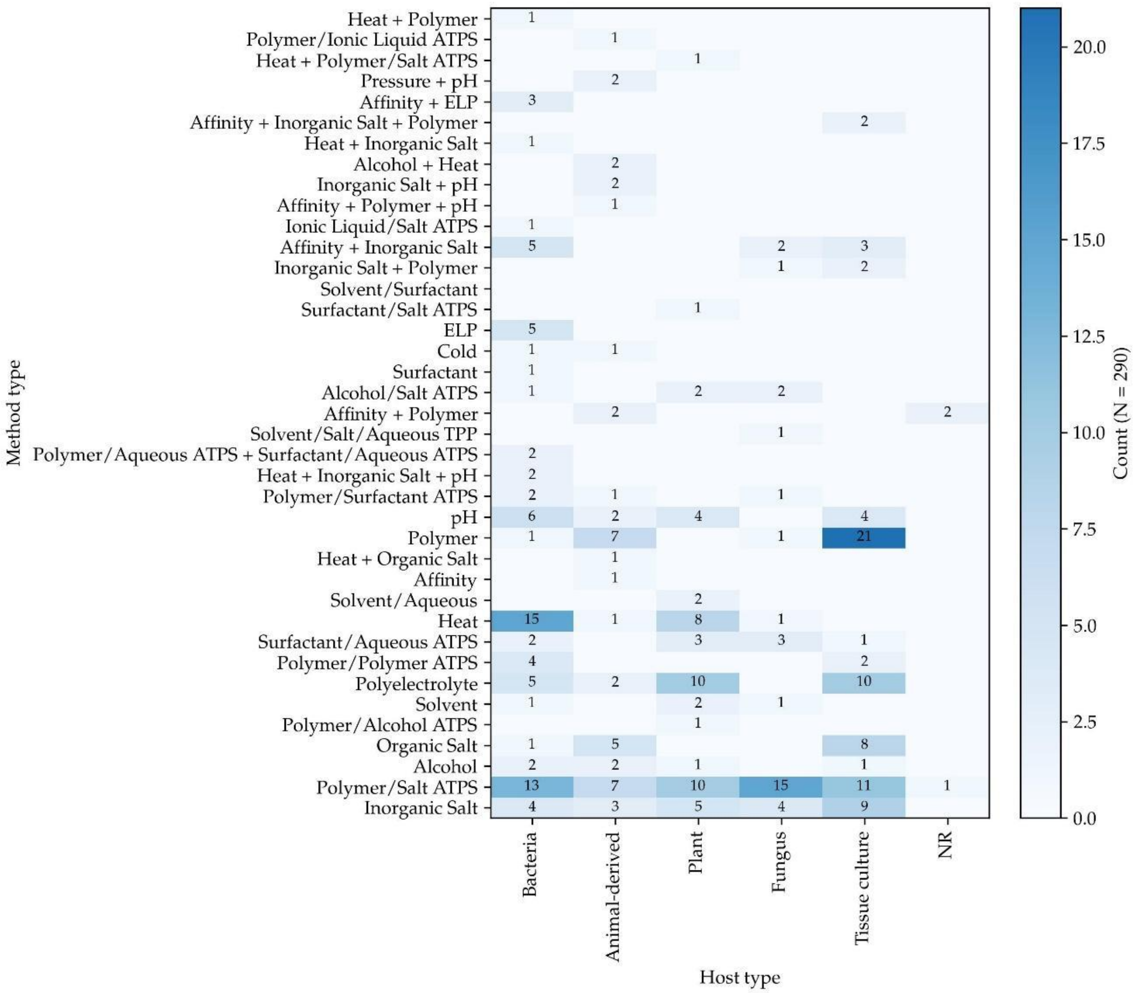


**Figure S7: Usage of different phase separation methods as a function of host type.** NR = not reported. Method types are precipitations unless ending with ATPS (aqueous two-phase separation) or TPP (three-phase partitioning).

### 2.2. Quantification of performance of phase separation protein purification methods

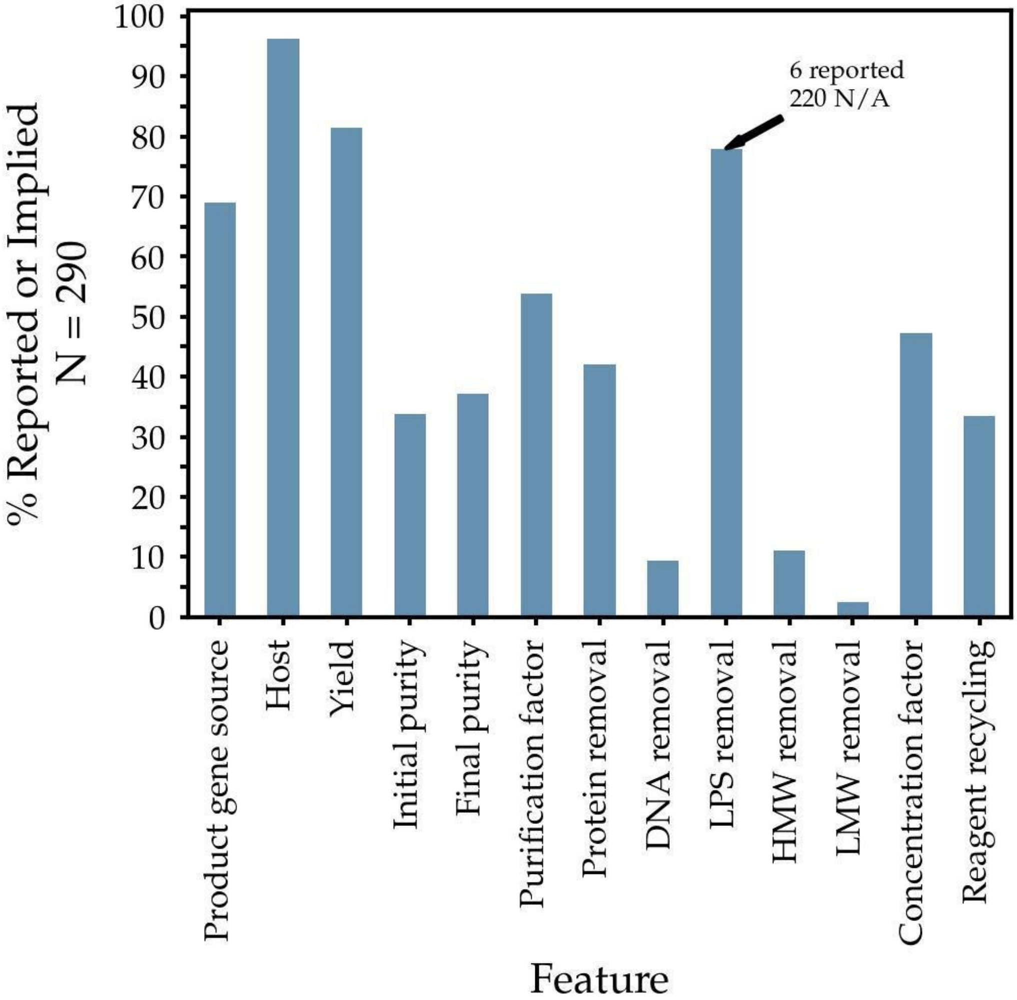


**Figure S8: Reporting frequency of method performance descriptors.** Many basic descriptions of purification performance or methodology needed for a complete comparative evaluation of protein purification methods are only sparsely reported. LPS: lipopolysaccharide (i.e., endotoxin). LMW: low molecular weight variants of the product. HMW: high molecular weight variants of the product. Methods listed are precipitations unless ending with ATPS (aqueous two-phase separation). For definitions of other terms, refer to Supplemental Methods. Descriptions were considered to be "implied" if they were either not applicable (e.g., LPS removal in non-bacterial hosts) or represented the most likely scenario based on the information provided (e.g., that a product gene was homologous to the host if nothing suggested otherwise).

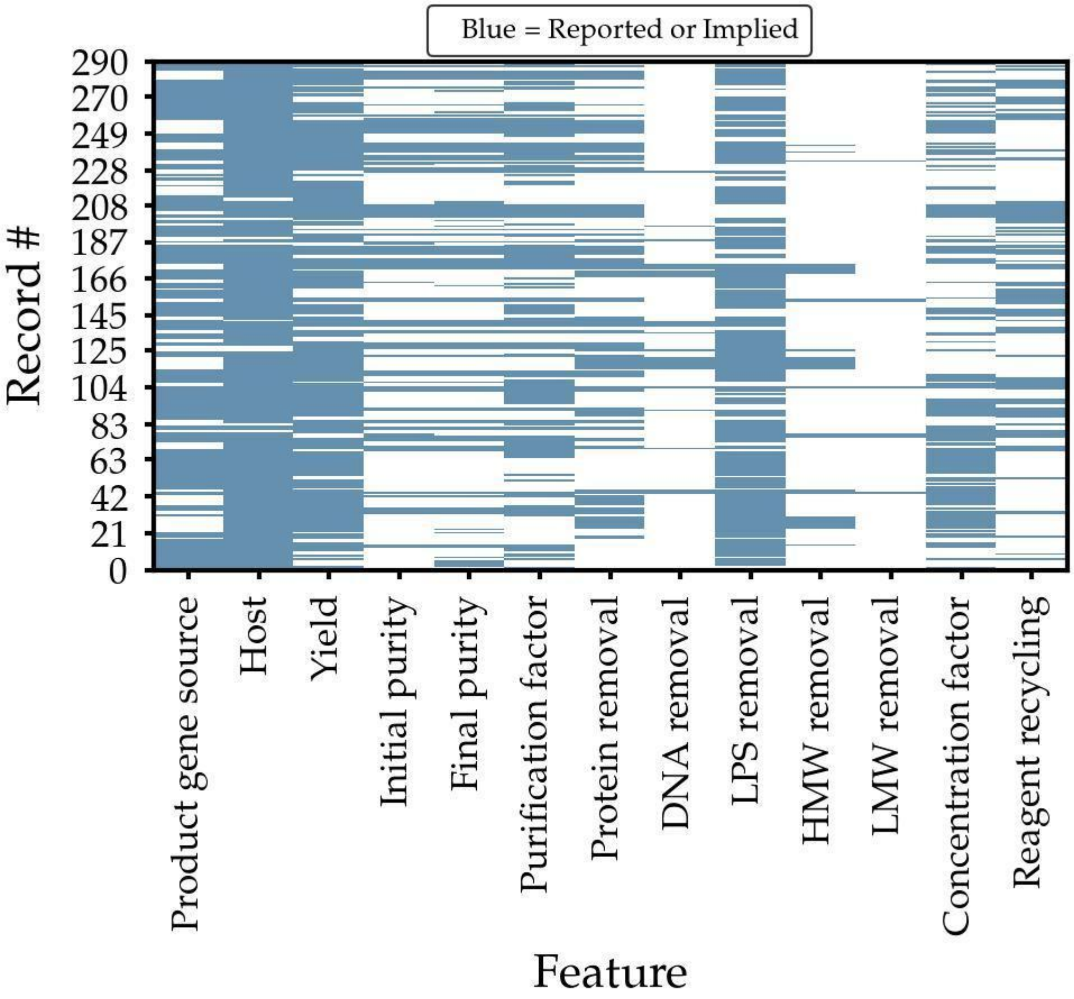


**Figure S9: Descriptor reporting completeness per record in the dataset.** Virtually all records lack some information required for a complete comparative evaluation of protein purification methods. LPS: lipopolysaccharide (i.e., endotoxin). LMW: low molecular weight variants of the product. HMW: high molecular weight variants of the product. Methods listed are precipitations unless ending with ATPS (aqueous two-phase separation). For definitions of other terms, refer to Supplemental Methods

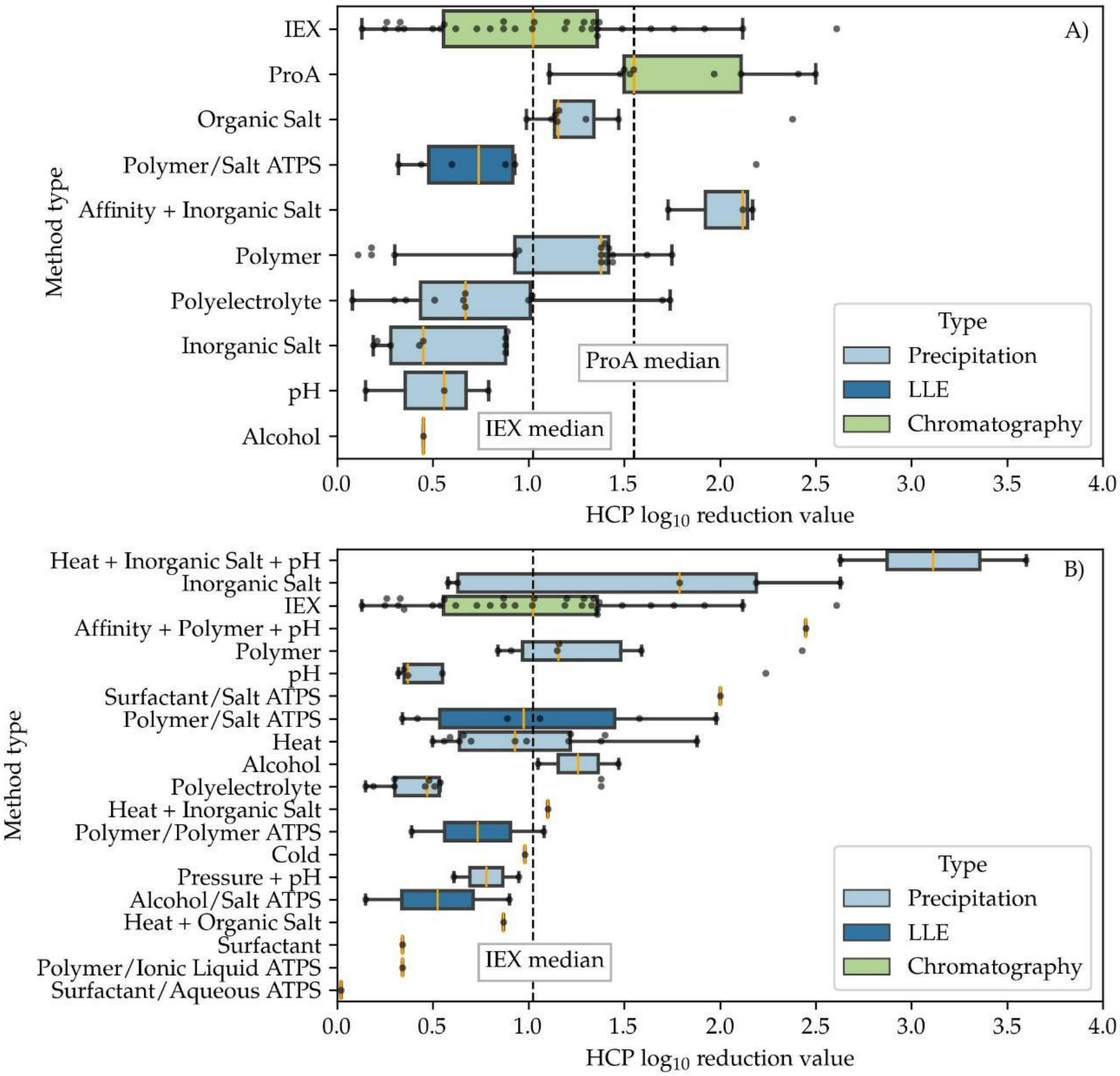


**Figure S10: Host cell protein removal by method type and product type.** Shown are values for phase separation-based methods and two common types of chromatography. A) Monoclonal antibodies purified by phase separation methods. B) Non-antibody products purified by phase separation methods. Box and whisker plots show the minimum, 25$^{th}$ percentile, 75$^{th}$ percentile and maximum values, respectively. Individual datapoints are shown as black circles. The median of each group is highlighted in orange.

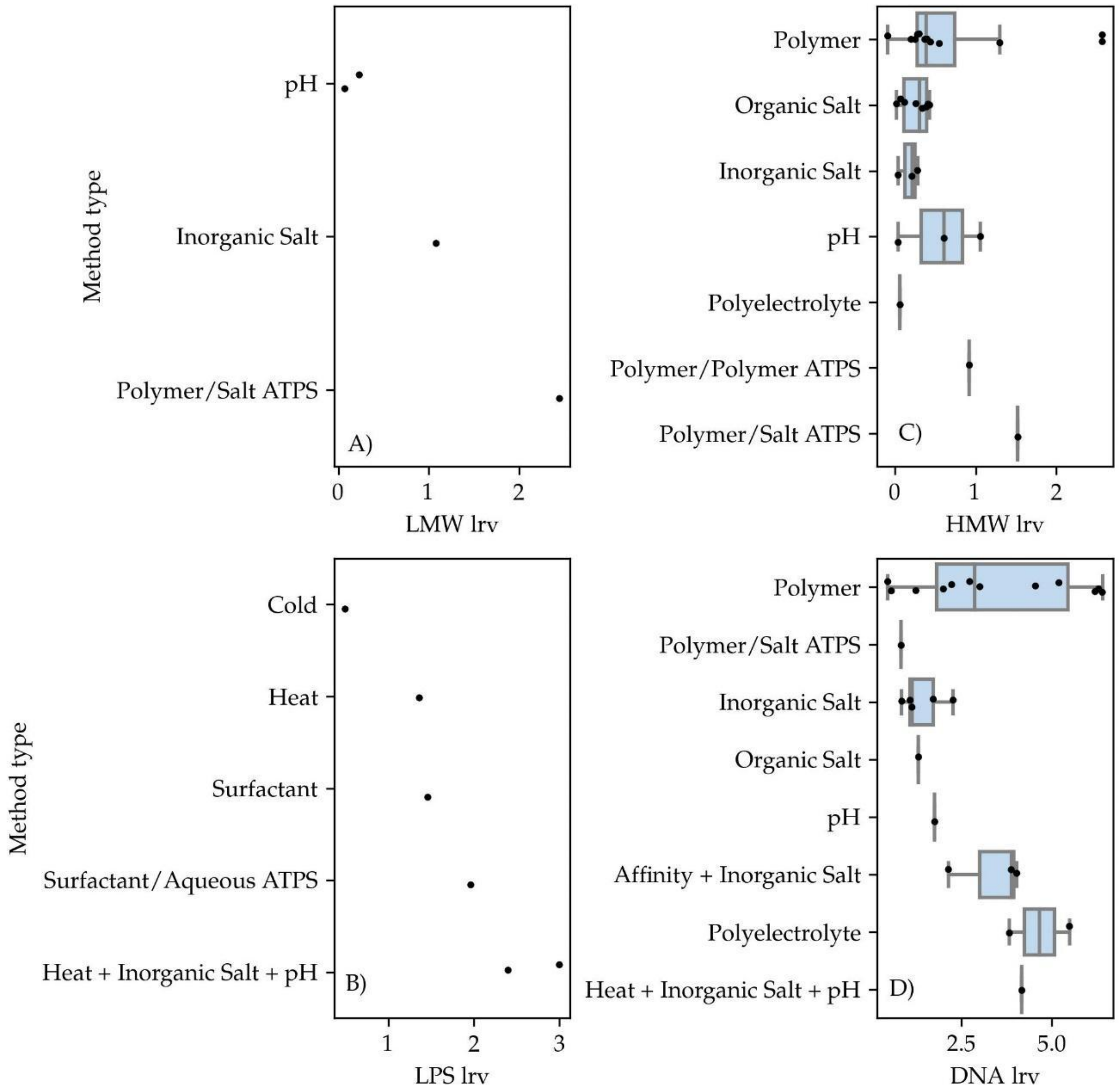


**Figure S11: Removal of LPS, DNA, LMW and HMW by phase separation.** Phase separation-based protein purification methods are capable of multiple order of magnitude reductions in DNA, low molecular weight, high molecular weight, and lipopolysaccharide contaminants. Box and whisker plots show minimum, 25$^{th}$ percentile, 75$^{th}$ percentile, and maximum values. LRV: $\log_{10}$ reduction value in contaminant mass. LPS: lipopolysaccharide (i.e., endotoxin). LMW: low molecular weight variants of the product. HMW: high molecular weight variants of the product. Methods listed are precipitations unless ending with ATPS (aqueous two-phase separation).

#### 2.3 Techno-economic analysis

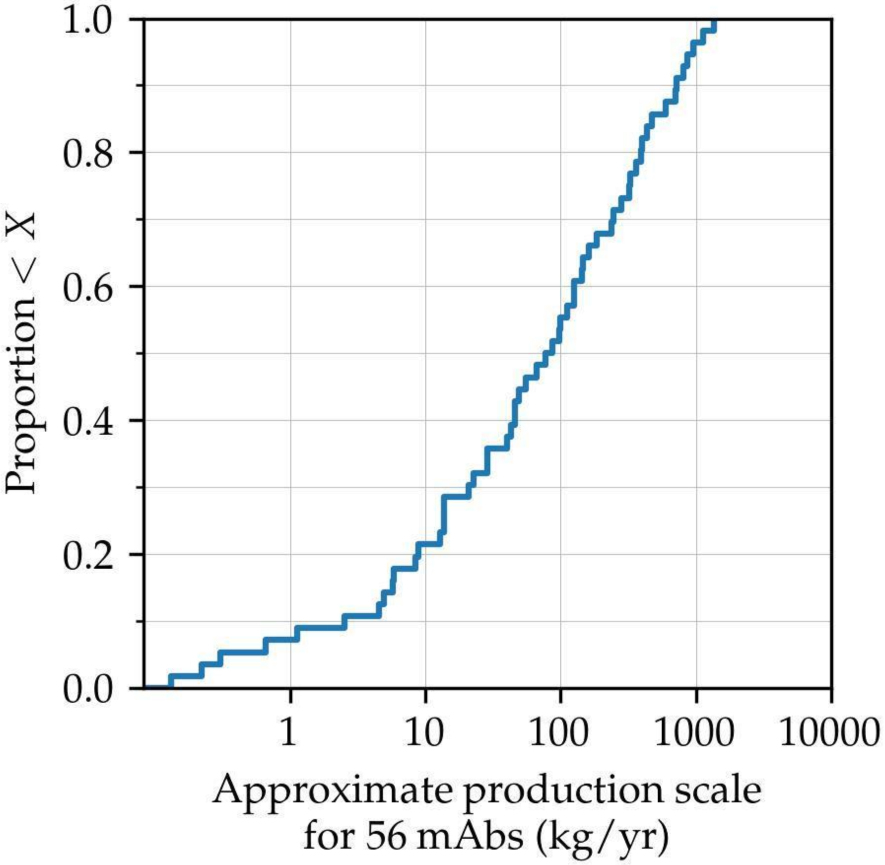


**Figure S12: Cumulative distribution of annual production scales for monoclonal antibody drug processes.** Adapted from data in Table S5.

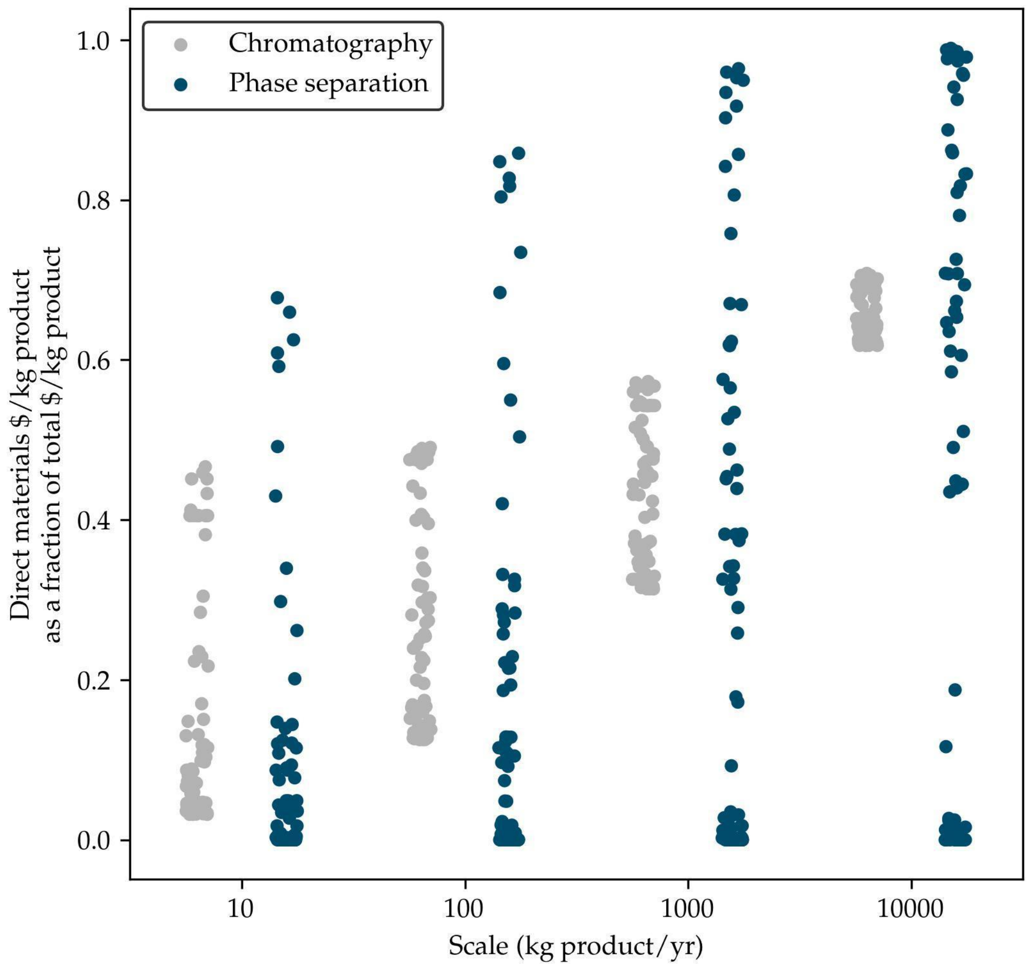


**Figure S13: Contribution of direct materials costs to total cost.** Direct material costs make up an increasing fraction of total costs per kg of product purified as production scale increases, for both phase separation methods and their ion-exchange chromatography-based alternatives. Direct materials: either phase-forming materials, in the case of phase-separation methods, or resin and chromatography buffers, for an alternative ion-exchange chromatography operation.

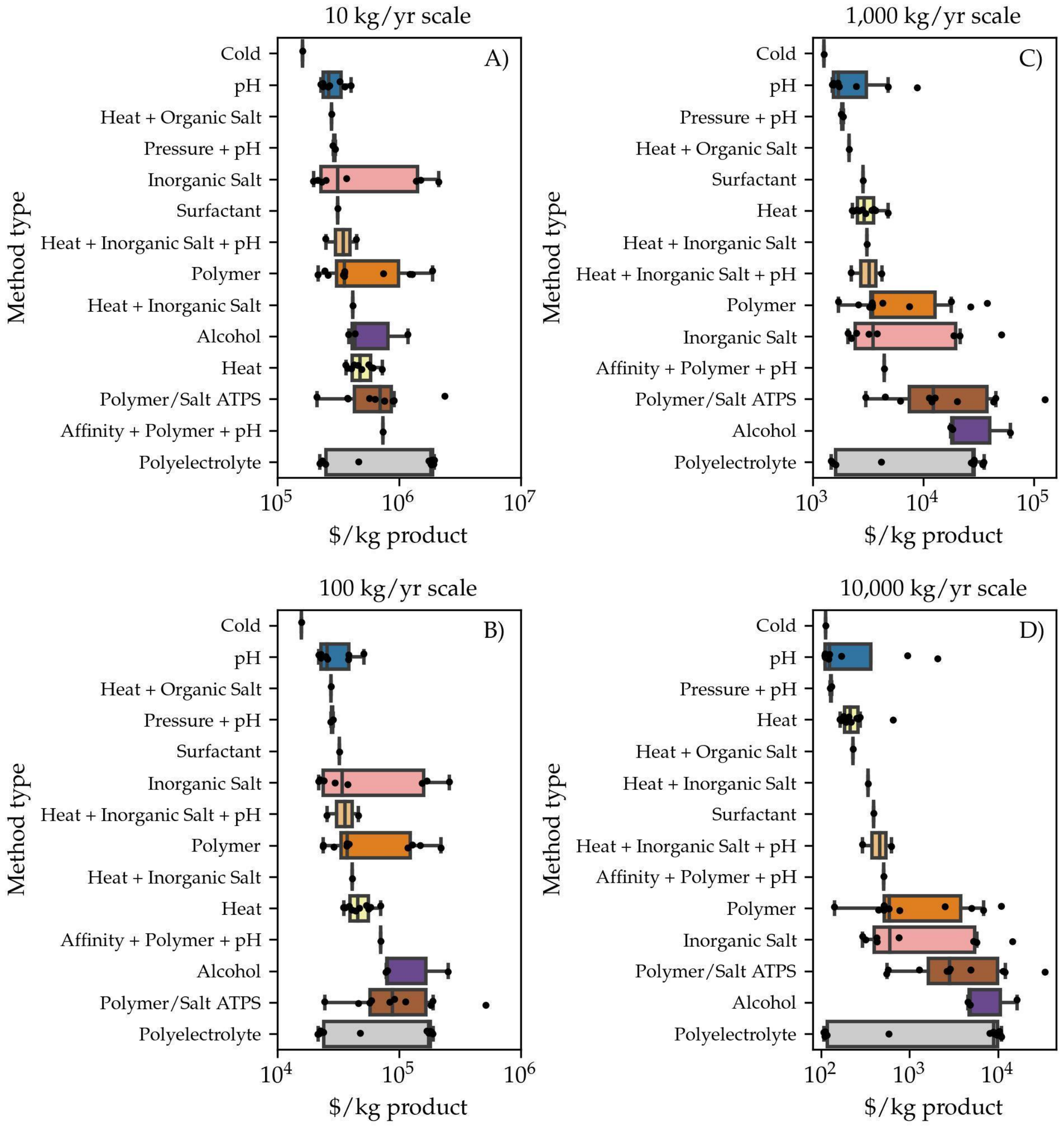


**Figure S14: Cost per kg of purified product per phase separation method.** Each panel shows a different level of annual product throughput. Box and whiskers show minimum, 25$^{th}$ percentile, 75$^{th}$ percentile, and maximum values. Methods are precipitations unless ending with ATPS (aqueous two-phase separation). Methods observed in the dataset but not shown did not report sufficient information for techno-economic analysis. Methods are sorted in ascending order of maximum cost in each panel, but box colors for each method are consistent among panels to allow comparison across production scales.

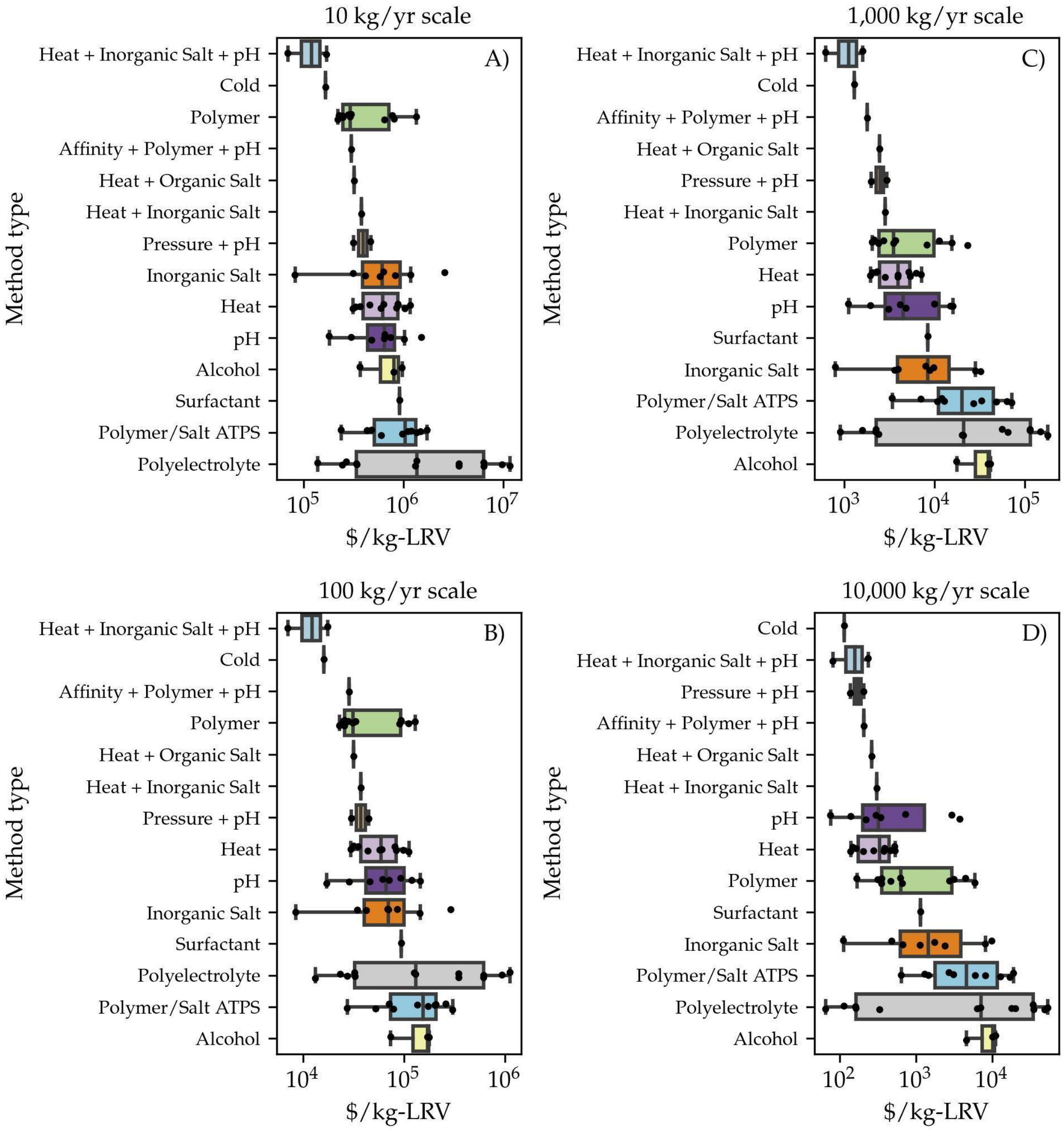


**Figure S15: Cost per kg of purified product per $\log_{10}$ reduction in host cell protein per phase separation method.** Each panel shows a different level of annual product throughput. Box and whiskers show minimum, $25^{th}$ percentile, $75^{th}$ percentile, and maximum values. Methods are precipitations unless ending with ATPS (aqueous two-phase separation). Methods observed in the dataset but not shown did not report sufficient information for techno-economic analysis. LRV: $\log_{10}$ reduction value, in this case, for host cell protein contaminants. Box colors for each method are consistent among panels to allow comparison across production scales.

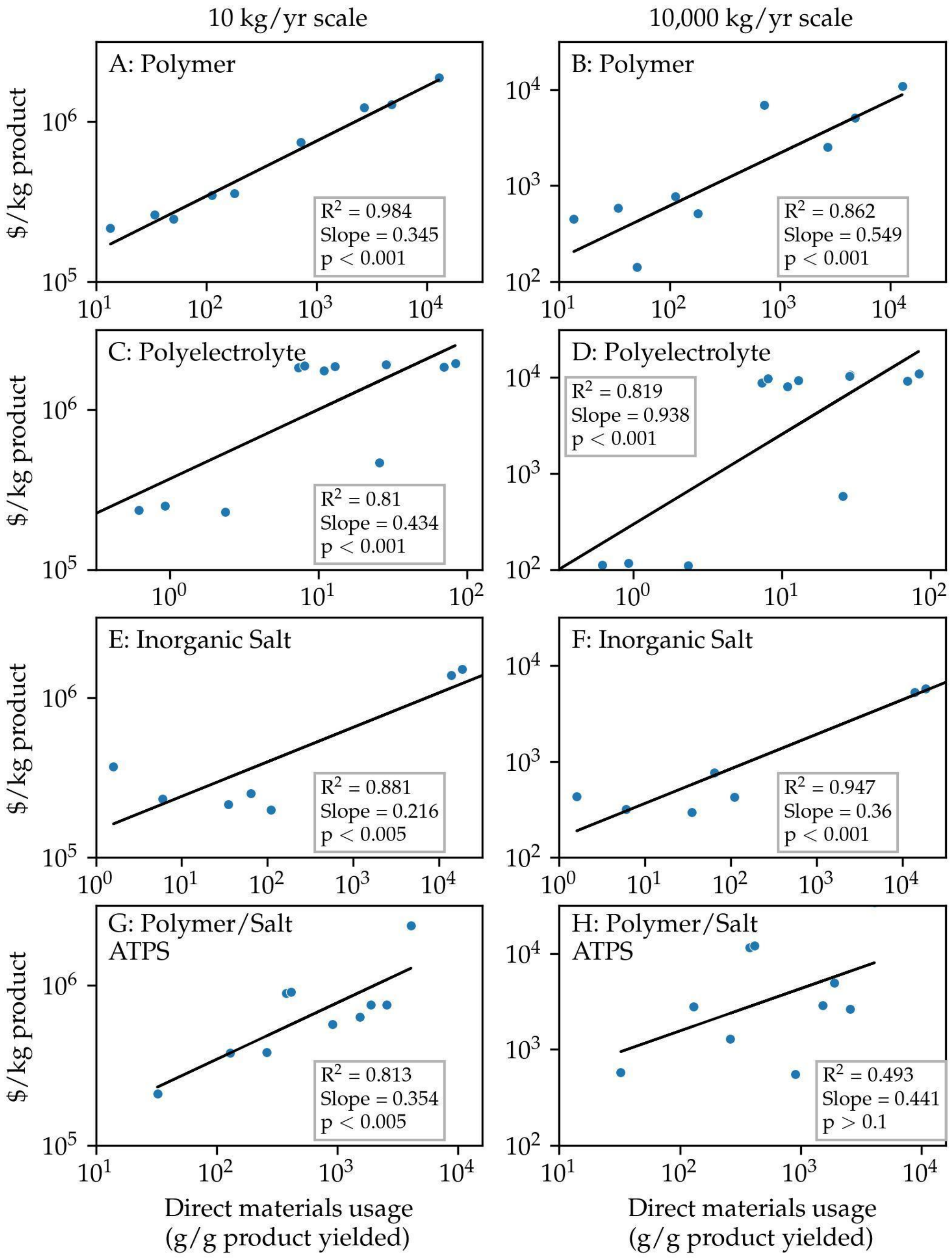


**Figure S16: Cost of product as a function of direct material usage rate.** Within a given phase separation method, total cost per kg of product purified is strongly predicted by the ratio of phase-forming materials mass to initial product mass. Black lines and inset boxes show the line of best fit and parameters for a linear regression of the log-scale data. Methods are precipitations unless ending with ATPS (aqueous two-phase separation). Left and right panels in each column show results at low and high levels of annual product throughput, respectively.

### 2.4. Implementation readiness

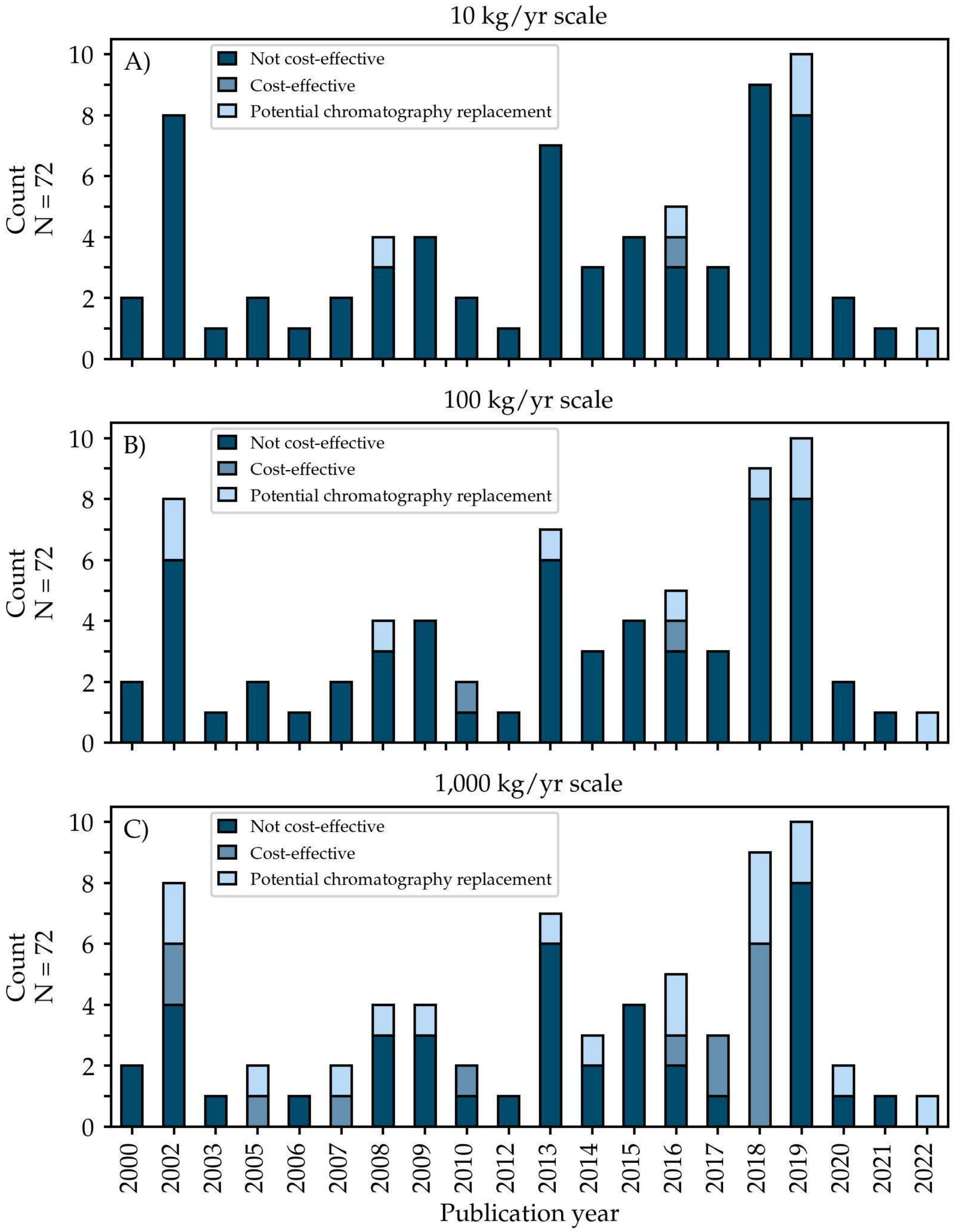


**Figure S17: Implementation readiness by year and annual product throughput.** For further information on the definitions of implementation readiness categories, refer to the text.

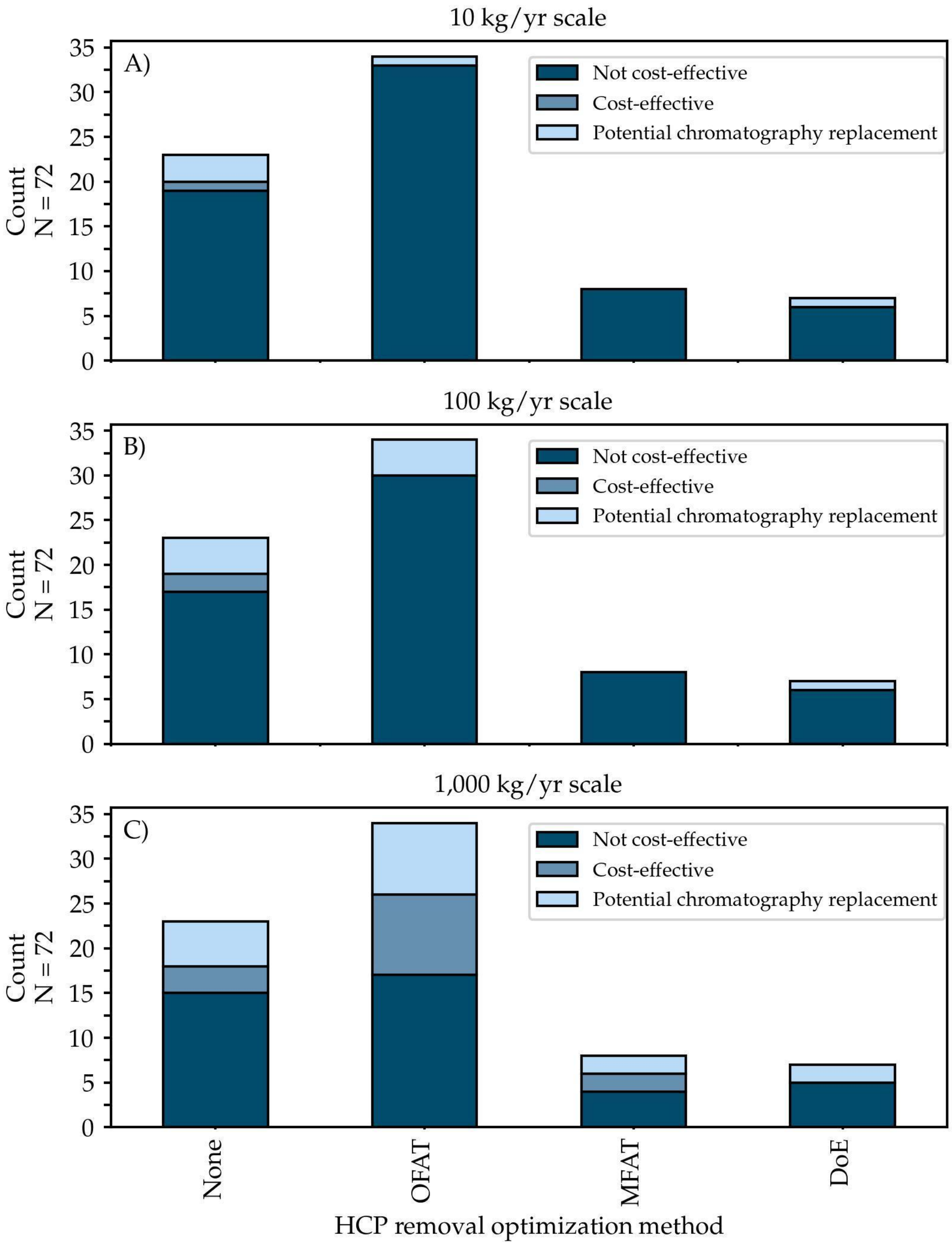


**Figure S18: Implementation readiness by method used to optimize host cell protein removal and of annual product throughput**. None: no optimization of host cell protein removal was reported. OFAT: one-factor-at-a-time optimization. MFAT: multiple-factors-at-a-time optimization. DoE: design of experiments. For further information on the definition of optimization methods and implementation readiness categories, refer to the text.

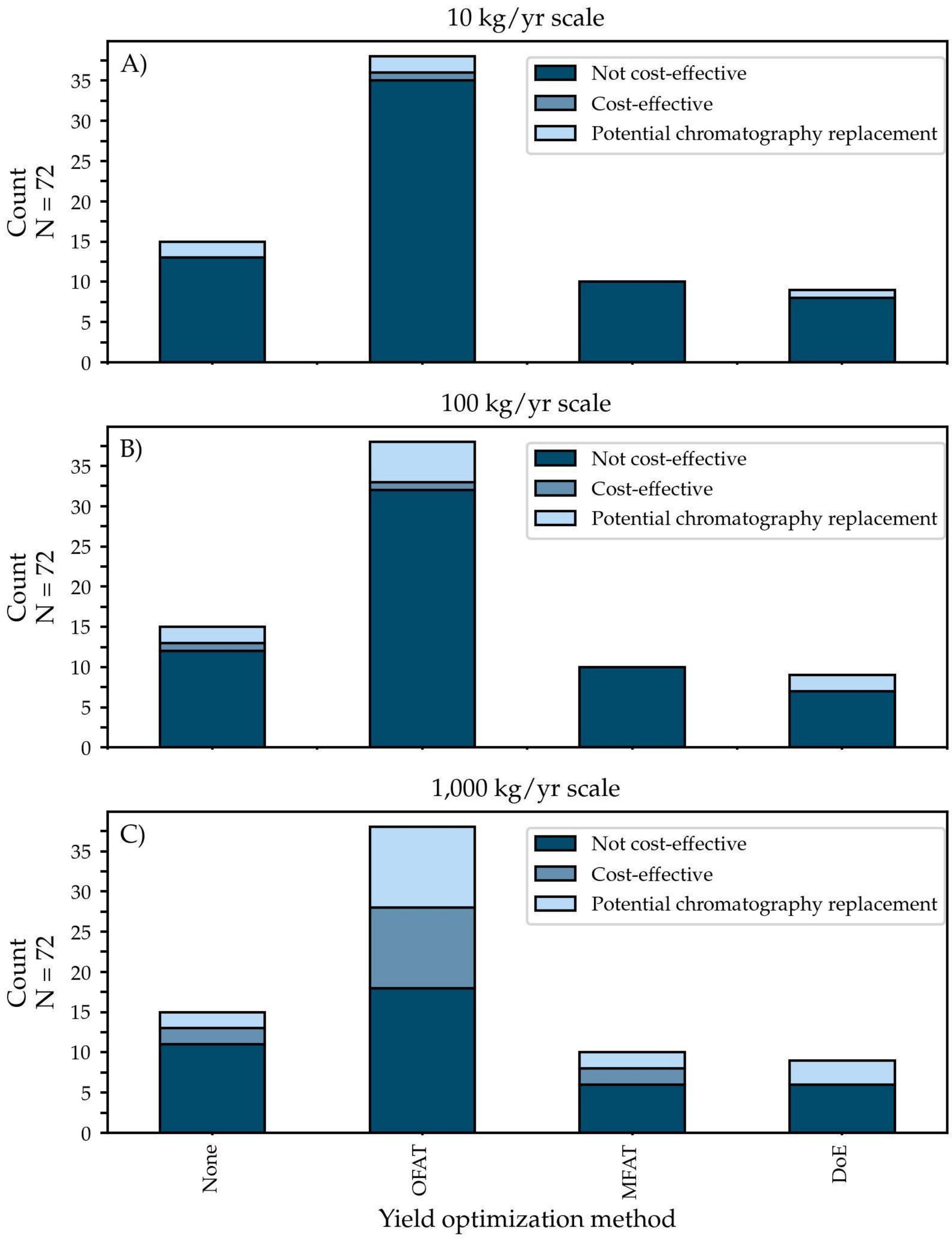


**Figure S19: Implementation readiness by method used to optimize product yield and of annual product throughput**. None: no optimization of yield was reported. OFAT: one-factor-at-a-time optimization. MFAT: multiple-factors-at-a-time optimization. DoE: design of experiments. For further information on the definition of optimization methods and implementation readiness categories, refer to the text.

### 2.5. Chromatography datasets

**Table S2: Typical product yields and host cell protein $\log_{10}$ reduction values for ion-exchange chromatography.** HCP: host cell protein. LRV: $\log_{10}$ reduction value in absolute mass of contaminant. For the data from Shukla et al., no yield was reported. However, because this study represented results in a pharmaceutical mAb bioprocess, we assumed a typical yield of 95% for that case (Kelley, 2007).

| **Source** | **Yield (%)** | **Initial purity (%)** | **Final purity (%)** | **HCP LRV** |
|---|---|---|---|---|
| (Hou et al., 2005)(Hou et al., 2005) | 63.87 | 35.40 | 95.30 | 1.76 |
| (Yu et al., 2021)(Yu et al., 2021) | 79.00 | 22.00 | 83.00 | 1.34 |
| (Guo et al., 2021)(Guo et al., 2021) | 67.03 | 45.00 | 82.27 | 0.93 |
| (Dux et al., 2006)(Dux et al., 2006) | 90.00 | 3.00 | 70.00 | 1.92 |
| (Dux et al., 2006)(Dux et al., 2006) | 76.00 | 70.00 | 80.00 | 0.35 |
| (Dux et al., 2006)(Dux et al., 2006) | 53.00 | 80.00 | 92.00 | 0.73 |
| (Bocanegra-Jiménez et al., 2021).*(Bocanegra-Jiménez et al., 2021)* | 79.17 | 55.00 | 86.00 | 0.80 |
| (Shukla et al., 2017)(Shukla et al., 2017) | 95.00 | 98.05 | 99.93 | 1.49 |
| (Shukla et al., 2017)(Shukla et al., 2017) | 95.00 | 99.16 | 99.95 | 1.29 |
| (Shukla et al., 2017)(Shukla et al., 2017) | 95.00 | 99.26 | 99.98 | 1.64 |
| (Shukla et al., 2017)(Shukla et al., 2017) | 95.00 | 99.30 | 99.97 | 1.37 |
| (Shukla et al., 2017)(Shukla et al., 2017) | 95.00 | 99.72 | 99.99 | 1.33 |
| (Shukla et al., 2017)(Shukla et al., 2017) | 95.00 | 99.80 | 99.99 | 1.36 |
| (Shukla et al., 2017)(Shukla et al., 2017) | 95.00 | 99.90 | 100.00 | 1.36 |

| | | | | |
|---|---|---|---|---|
| (Shukla et al., 2017)(Shukla et al., 2017) | 95.00 | 99.92 | 100.00 | 1.28 |
| (Shukla et al., 2017)(Shukla et al., 2017) | 95.00 | 99.93 | 100.00 | 1.20 |
| (Shukla et al., 2017)(Shukla et al., 2017) | 95.00 | 99.95 | 100.00 | 1.02 |
| (Shukla et al., 2017)(Shukla et al., 2017) | 95.00 | 99.98 | 100.00 | 0.62 |
| (Shukla et al., 2017)(Shukla et al., 2017) | 95.00 | 99.97 | 100.00 | 0.87 |
| (Shukla et al., 2017)(Shukla et al., 2017) | 95.00 | 99.99 | 100.00 | 0.50 |
| (Shukla et al., 2017)(Shukla et al., 2017) | 95.00 | 99.99 | 100.00 | 0.32 |
| (Olsen et al., 2013)(Olsen et al., 2013) | 115.38 | 0.79 | 78.95 | 2.61 |
| (A. Li et al., 2021)(Li et al., 2021) | 79.49 | 71.00 | 78.00 | 0.26 |
| (Burkhardt et al., 2007)(Burkhardt et al., 2007) | 69.00 | 9.00 | 90.00 | 2.12 |
| (Freiherr von Roman et al., 2014)(Freiherr von Roman et al., 2014) | 97.00 | 92.00 | 96.00 | 0.33 |
| (Xiang et al., 2010)(Xiang et al., 2010) | 90.91 | 94.00 | 95.00 | 0.13 |
| (Xiang et al., 2010)(Xiang et al., 2010) | 70.00 | 95.00 | 99.00 | 0.87 |
| (C. Huang et al., 2005)(Huang et al., 2005) | 96.70 | 85.00 | 95.00 | 0.54 |
| (Hofmann et al., 2013)(Hofmann et al., 2013) | 97.32 | 35.00 | 85.00 | 1.03 |

| | | | | |
|---|---|---|---|---|
| (Hofmann et al., 2013)(Hofmann et al., 2013) | 91.74 | 85.00 | 95.00 | 0.56 |
| (Hofmann et al., 2013)(Hofmann et al., 2013) | 70.97 | 45.00 | 90.00 | 1.19 |
| (Hofmann et al., 2013)(Hofmann et al., 2013) | 82.95 | 90.00 | 93.00 | 0.25 |

**Table S3: Typical host cell protein $log_{10}$ reduction values for Protein A chromatography.** Data are from (Shukla et al., 2017). PPM: parts of contaminant per million parts of product. HCP: host cell protein. LRV: $log_{10}$ reduction value in absolute mass of contaminant. For calculating HCP LRV from ppms, a product yield of 95% was assumed as typical for Protein A in a pharmaceutical process. (Kelley, 2007).

| **Initial HCP ppm (assumed—see caption)** | **HCP ppm post-ProA** | **HCP LRV** |
|---|---|---|
| 241948.47 | 19936.71 | 1.11 |
| 241948.47 | 8453.89 | 1.48 |
| 241948.47 | 8137.43 | 1.50 |
| 241948.47 | 7504.52 | 1.53 |
| 241948.47 | 7097.65 | 1.55 |
| 241948.47 | 2757.69 | 1.97 |
| 241948.47 | 1989.15 | 2.11 |
| 241948.47 | 994.58 | 2.41 |
| 241948.47 | 813.74 | 2.50 |

**Table S4: Typical DNA $log_{10}$ reduction values for ion-exchange chromatography.** Data are from (Berthold and Walter, 1994). LRV: $log_{10}$ reduction value in absolute mass of contaminant.

| Fold reduction in DNA | DNA LRV |
|---|---|
| 2,360,000.00 | 6.37 |
| 1,500,000.00 | 6.18 |
| 1,320,000.00 | 6.12 |
| 1,500,000.00 | 6.18 |
| 1,050,000.00 | 6.02 |
| 1,546,000.00 | 6.19 |

**Table S5: Market data on therapeutic mAbs sold in the US in 2020.** Sales price data reflect the lowest prices listed in 2020 procurement contracts with the US Department of Veterans Affairs, and total revenues and manufacturer COGS data are from manufacturer annual financial reports from 2019 or 2018. Drug mass sold is estimated simply by dividing total revenues by per-mass price. COGS per gram of drug is estimated by multiplying the per-gram sales price by the manufacturer's total COGS as a fraction of total revenues.

| mAb [Manufacturer] | Lowest sales price ($/g) | Global mAb revenues ($M/yr) | Manufacturer COGS (% of total revenue) | Estimated drug mass sold (kg) | Estimated COGS ($/g) |
|---|---|---|---|---|---|
| Adalimumab [Abbvie] | $31,264 | $14,864 | 18.18 | 475.43 | $5,682.44 |
| Risankizumab [Abbvie] | $72,380 | $355 | 18.18 | 4.90 | $13,155.31 |
| Eculizumab [Alexion Pharmaceuticals] | $16,608 | $3,946 | 7.01 | 237.62 | $1,164.23 |
| Ravulizumab- Cwvz [Alexion Pharmaceuticals] | $16,305 | $339 | 7.01 | 20.79 | $1,142.99 |
| Erenumab-Aooe [Amgen] | $3,062 | $306 | 9.07 | 99.92 | $277.78 |
| Evolocumab [Amgen] | $1,108 | $661 | 9.07 | 596.44 | $100.52 |
| Panitumumab [Amgen] | $8,508 | $744 | 9.07 | 87.45 | $771.70 |
| Romosozumab- Aqqg [Amgen] | $6,547 | $189 | 9.07 | 28.87 | $593.88 |
| Trastuzumab- Anns [Amgen] | $5,597 | $226 | 9.07 | 40.38 | $507.66 |
| Benralizumab [Astrazeneca] | $120,425 | $686 | 16.74 | 5.70 | $20,157.08 |
| Durvalumab [Astrazeneca] | $5,242 | $1,469 | 16.74 | 280.24 | $877.42 |
| Palivizumab [Astrazeneca] | $20,849 | $269 | 16.74 | 12.90 | $3,489.69 |
| Brodalumab [Bausch Health] | $6,235 | $28 | 25.97 | 4.49 | $1,618.92 |
| Natalizumab [Biogen] | $13,119 | $1,892 | 12.33 | 144.22 | $1,618.18 |
| Elotuzumab [Bristol-Myers Squibb] | $4,615 | $357 | 25.72 | 77.35 | $1,187.09 |
| Ipilimumab [Bristol-Myers Squibb] | $108,709 | $1,489 | 25.72 | 13.70 | $27,961.44 |
| Nivolumab [Bristol-Myers Squibb] | $18,188 | $7,204 | 25.72 | 396.10 | $4,678.09 |
| Cetuximab [Eli Lilly] | $4,318 | $635 | 20.19 | 147.14 | $871.92 |
| Galcanezumab- Gnlm [Eli Lilly] | $3,561 | $163 | 20.19 | 45.63 | $719.11 |
| Ixekizumab [Eli Lilly] | $48,053 | $1,366 | 20.19 | 28.44 | $9,703.97 |

| | | | | | |
|---|---|---|---|---|---|
| Necitumumab [Eli Lilly] | $4,099 | $10 | 20.19 | 2.51 | $827.81 |
| Olaratumab [Eli Lilly] | $3,694 | $203 | 20.19 | 54.95 | $746.06 |
| Ramucirumab [Eli Lilly] | $8,324 | $925 | 20.19 | 111.14 | $1,680.96 |
| Avelumab [EMD Serono] | $5,814 | $79 | 21.68 | 13.58 | $1,260.69 |
| Ado-Trastuzumab Emtansine [Genentech] | $22,169 | $1,483 | 22.83 | 66.88 | $5,061.62 |
| Atezolizumab [Genentech] | $5,553 | $1,996 | 22.83 | 359.37 | $1,267.86 |
| Bevacizumab [Genentech] | $5,483 | $7,528 | 22.83 | 1372.97 | $1,251.87 |
| Emicizumab- Kxwh [Genentech] | $62,444 | $1,424 | 22.83 | 22.80 | $14,257.47 |
| Obinutuzumab [Genentech] | $4,646 | $587 | 22.83 | 126.43 | $1,060.81 |
| Ocrelizumab [Genentech] | $40,740 | $3,946 | 22.83 | 96.85 | $9,301.98 |
| Omalizumab [Genentech] | $5,256 | $2,095 | 22.83 | 398.62 | $1,200.17 |
| Pertuzumab [Genentech] | $8,632 | $3,748 | 22.83 | 434.21 | $1,970.96 |
| Polatuzumab Vedotin-Piiq [Genentech] | $81,838 | $54 | 22.83 | 0.66 | $18,685.59 |
| Ranibizumab [Genentech] | $1,715,533 | $1,943 | 22.83 | 1.13 | $391,698.64 |
| Rituximab [Genentech] | $6,089 | $6,892 | 22.83 | 1131.88 | $1,390.28 |
| Tocilizumab [Genentech] | $3,478 | $2,459 | 22.83 | 707.09 | $794.06 |
| Trastuzumab [Genentech] | $7,402 | $6,426 | 22.83 | 868.17 | $1,690.00 |
| Alemtuzumab [Genzyme] | $1,361,914 | $293 | 28.76 | 0.22 | $391,621.47 |
| Caplacizumab- Yhdp [Genzyme] | $494,383 | $64 | 28.76 | 0.13 | $142,160.86 |
| Belimumab [GlaxoSmithKline] | $3,152 | $775 | 25.66 | 246.00 | $808.92 |
| Mepolizumab [GlaxoSmithKline] | $21,260 | $972 | 25.66 | 45.70 | $5,455.68 |
| Daratumumab [Johnson & Johnson] | $3,135 | $2,998 | 24.45 | 956.18 | $766.66 |
| Golimumab [Johnson & Johnson] | $17,556 | $2,188 | 24.45 | 124.63 | $4,292.66 |

| | | | | | |
|---|---|---|---|---|---|
| Guselkumab [Johnson & Johnson] | $74,264 | $1,012 | 24.45 | 13.63 | $18,158.89 |
| Infliximab [Johnson & Johnson] | $5,451 | $4,380 | 24.45 | 803.60 | $1,332.74 |
| Ustekinumab [Johnson & Johnson] | $8,876 | $6,361 | 24.45 | 716.63 | $2,170.41 |
| Pembrolizumab [Merck] | $34,580 | $11,084 | 21.68 | 320.54 | $7,498.19 |
| Canakinumab [Novartis] | $79,994 | $671 | 24.29 | 8.39 | $19,431.31 |
| Secukinumab [Novartis] | $21,945 | $3,551 | 24.29 | 161.82 | $5,330.53 |
| Cemiplimab-Rwic [Regeneron] | $19,769 | $176 | 59.64 | 8.90 | $11,789.98 |
| Alirocumab [Sanofi Aventis] | $1,462 | $271 | 28.76 | 185.35 | $420.49 |
| Dupilumab [Sanofi Aventis] | $7,195 | $2,339 | 28.76 | 325.09 | $2,068.88 |
| Sarilumab [Sanofi Aventis] | $4,844 | $209 | 28.76 | 43.20 | $1,392.83 |
| Brentuximab Vedotin [Seattle Genetics] | $108,120 | $628 | 6.75 | 5.81 | $7,292.97 |
| Fremanezumab- Vfrm [Teva Pharmaceuticals] | $1,909 | $93 | 47.09 | 48.72 | $898.88 |
| Dinutuximab [United Therapeutics] | $377,992 | $114 | 10.00 | 0.30 | $37,803.84 |

**2.6. Records used for techno-economic analysis.** Phase separation records from the dataset that had sufficient information to perform a techno-economic analysis. For references that included more than one record, additional information is provided to identify the specific method.

| Reference | Additional information | Method type | Success at 100 kg/yr | Success at 1000 kg/yr | Success at 10000 kg/yr |
|---|---|---|---|---|---|
| Cabrera-Crespo, J. et. al. (2000). Biotechnol. Appl. Biochem. 31, 101–106. | 25% ethanol and 0.6% chloroform, 10 °C, pH 7.2, overnight | Alcohol | Not cost effective | Not cost effective | Not cost effective |
| Cabrera-Crespo, J. et. al. (2000). Biotechnol. Appl. Biochem. 31, 101–106. | 2.3 g of sodium octanoate per 100 g protein, 2 mM EDTA, pH 6.5, 70 °C, 1 h | Heat/Organic Salt | Not cost effective | Not cost effective | Cost effective |
| Capito, F., et. al. (2014). Biotechnol. J. 9, 698–701. | - | Polyelectrolyte | Not cost effective | Not cost effective | Potential replacement |
| Chen, Q., et. al. (2016). J. Biotechnol. 236, 128–140. | - | Inorganic Salt | Not cost effective | Not cost effective | Not cost effective |

| | | | | | |
|---|---|---|---|---|---|
| Decker, J. S., et.al. (2020). Front Bioeng Biotechnol 8, 1020. | - | Heat/Inorganic Salt/pH | Not cost effective | Potential replacement | Potential replacement |
| Decker, J. S., et. al. (2022). Biochem. Eng. J. 181, 108403 | - | Heat/Inorganic Salt/pH | Potential replacement | Potential replacement | Potential replacement |
| Diederich, P., et. al. (2015). Biotechnol. Prog. 31, 957–973. | 32% w/w PEG 600 and 1% w/w potassium phosphate | Polymer | Not cost effective | Not cost effective | Not cost effective |
| Diederich, P., et. al. (2015). Biotechnol. Prog. 31, 957–973. | Same as above plus 24.4% PEG600, 11.5% ammonium sulfate and 3% NaCl (w/w) | Polymer/Salt ATPS | Not cost effective | Not cost effective | Not cost effective |
| DiScipio, R. G., et. al. (2016). Protein Expr. Purif. 121, 118–124. | 4% w/v PEG-4000, 46.6 g/L total protein with 17 mg/L | Polymer | Cost effective | Cost effective | Cost effective |
| DiScipio, R. G., et. al. (2016). Protein Expr. Purif. 121, 118–124. | 35% saturated ammonium sulfate, 21.6 g/L total protein with 17 mg/L fibulin | Inorganic Salt | Potential replacement | Potential replacement | Potential replacement |
| Dutra G., et. al. (2020). Eng. J. 20, 265-274 | - | Inorganic Salt | Not cost effective | Not cost effective | Cost effective |
| Eguia F. et. al. (2021). Appl. Microbiol. Biotechnol. 105, 169-183 | - | pH | Not cost effective | Not cost effective | Cost effective |
| Figueiredo D. B., et. al. (2017) Appl. Microbiol. Biotechnol. 101, 2305-2317 | 0.1% cetyltrimethylammonium bromide, pH 6.5 | Polyelectrolyte/ Surfactant | Not cost effective | Not cost effective | Not cost effective |
| Figueiredo D. B., et. al. (2017) Appl. Microbiol. Biotechnol. 101, 2305-2317 | -20 °C, pH 4.0, 24 h | Cold | Not cost effective | Cost effective | Cost effective |
| Geng F., et. al. (2013) J. Sep. Sci. 36, 3717-3722 | 5% w/v PEG-8000, treat with 9% w/v PEG-8000 | Polymer | Potential replacement | Potential replacement | Potential replacement |
| Geng F., et. al. (2019) Poult. Sci. 98, 1545-1550 | 15% w/v PEG-8000, 30 min, 10 °C, pH 6.5, 100 mM NaCl | Polymer | Not cost effective | Not cost effective | Not cost effective |
| Geng F., et. al. (2019) Poult. Sci. 98, 1545-1550 | pH 4.5, 4 C, 15 min | pH | Not cost effective | Not cost effective | Cost effective |
| Golubovic M., et. al. (2005) J. Agric. Food Chem. 53, 5265-5269 | - | pH | Not cost effective | Potential replacement | Potential replacement |
| Großhans S., et. al. (2019) Bioprocess Biosyst Eng. 42, 1039-1051 | 8 mg/mL mAb, 0.5 mg/mL HCP in untreated mock CCF, 10% w/w PEG 6000, pH 7.5, 50 mM Tris, 30 min, 22 °C | Polymer | Not cost effective | Not cost effective | Not cost effective |
| Großhans S., et. al. (2019) Bioprocess Biosyst Eng. 42, 1039-1051 | 4 mg/mL mAb, 0.5 mg/mL HCP in untreated mock CCF, 12.73% w/w PEG 6000, pH 7.5, 50 mM Tris, 30 min, 22 °C | Polymer | Not cost effective | Not cost effective | Not cost effective |
| Gu Z. & Glatz C. E. (2007) J. Chromatogr. B Analyt. Technol. Biomed. Life Sci. 845, 38-50 | 15 ug product, 0.1 g corn endosperm, 15.7% w/w PEG 3350, 8.9% w/w Na2SO4, 4.5% | Polymer/Salt ATPS | Not cost effective | Cost effective | Cost effective |

| | | | | | |
|---|---|---|---|---|---|
| | w/w NaCl, Vr = 0.71, product to top, 1 hr, 22 °C | | | | |
| Gu Z. & Glatz C. E. (2007) J. Chromatogr. B Analyt. Technol. Biomed. Life Sci. 845, 38-50 | 0.01 g corn germ and 15 ug product per g ATPS. 16.3% w/w PEG 1450, 13.5% w/w, 8.5% w/w NaCl, Vr = 0.67, product to bottom, 1 h, 22 °C | Polymer/Salt ATPS | Not cost effective | Potential replacement | Potential replacement |
| Hammerschmidt N., et. al. (2015) Biotechnol. J. 10, 1196-1205 | 250 mM CaCl2, pH 8.0 | Inorganic Salt | Not cost effective | Not cost effective | Not cost effective |
| Hammerschmidt N., et. al. (2015) Biotechnol. J. 10, 1196-1205 | 25% v/v ethanol, -10 °C | Alcohol | Not cost effective | Not cost effective | Not cost effective |
| Ishihara T., et. al. (2019) J. Chromatogr. B Analyt. Technol. Biomed. Life Sci. 1121, 72-81 | CCF, pH 4.5, 2.7 g/L mAb, 1 hr | pH | Not cost effective | Not cost effective | Cost effective |
| Ishihara T., et. al. (2019) J. Chromatogr. B Analyt. Technol. Biomed. Life Sci. 1121, 72-81 | CCF, pH 4.5, 4.7 g/L mAb, 1 hr | pH | Not cost effective | Not cost effective | Not cost effective |
| Janoschek L., et. al. (2014) ) J. Chromatogr. B Analyt. Technol. Biomed. Life Sci. 965, 72-78 | - | Affinity/pH/Polymer | Not cost effective | Potential replacement | Potential replacement |
| Kateja N., et. al. (2016) Biotechnol. J. 11, 1320-1331 | - | pH | Not cost effective | Not cost effective | Cost effective |
| Kepka C., et. al. (2005) - J. Chromatogr. A. 1075, 33-41 | - | Polymer/Salt ATPS | Not cost effective | Cost effective | Cost effective |
| Kim H. J., et. al. (2010) Protein Expr. Purif. 70, 68-74 | - | Inorganic Salt | Cost effective | Cost effective | Cost effective |
| Kim C. K., et. al. (2013) PLoS One, 8(11):e80109 | - | pH | Not cost effective | Not cost effective | Not cost effective |
| Kim C. K. et. al. (2014) Appl. Biochem. Biotechnol. 172, 2425-2435 | - | pH | Not cost effective | Not cost effective | Not cost effective |
| Kim H. et. al. (2019) J. Virol. Methods. 269, 26-29 | 7.5% w/w PEG 8000 | Polymer | Potential replacement | Potential replacement | Potential replacement |
| Kim H. et. al. (2019) J. Virol. Methods. 269, 26-29 | 750% w/w ammonium sulfate | Inorganic Salt | Potential replacement | Potential replacement | Potential replacement |
| Ko K. Y. & Ahn D. U. (2008) Poult. Sci. 87, 1441-1450 | - | Alcohol | Not cost effective | Not cost effective | Not cost effective |
| Lan L. H., et. al. (2016) Biotechnol J. 11, 1595-1604 | - | Heat/Inorganic Salt | Not cost effective | Potential replacement | Potential replacement |

| | | | | | |
|---|---|---|---|---|---|
| Li Z., et. al. (2018) J. Chromatogr. A. 1581-82, 21-28 | For Hepatitis B core protein. Clarified lysate, 60 °C, 30 min. | Heat | Not cost effective | Cost effective | Cost effective |
| Li Z., et. al. (2018) J. Chromatogr. A. 1581-82, 21-28 | For Hepatitis B core protein. Supernatant, then 70 °C, 30 min | Heat | Not cost effective | Potential replacement | Potential replacement |
| Li Z., et. al. (2018) J. Chromatogr. A. 1581-82, 21-28 | For Ovalbumin-HBc. Clarified lysate, 60 °C, 30 min | Heat | Not cost effective | Cost effective | Cost effective |
| Li Z., et. al. (2018) J. Chromatogr. A. 1581-82, 21-28 | For Ovalbumin-HBc. Supernatant, then 70 °C, 30 min | Heat | Not cost effective | Cost effective | Cost effective |
| Li Z., et. al. (2018) J. Chromatogr. A. 1581-82, 21-28 | For Matrix 2 protein-HBc. Clarified lysate, 60 °C, 30 min. | Heat | Not cost effective | Cost effective | Cost effective |
| Li Z., et. al. (2018) J. Chromatogr. A. 1581-82, 21-28 | For Matrix 2 protein-HBc. Supernatant, then 70 °C, 30 min | Heat | Not cost effective | Potential replacement | Potential replacement |
| Li Z., et. al. (2018) J. Chromatogr. A. 1581-82, 21-28 | For Nuclear protein-HBc. Clarified lysate, 60 °C, 30 min | Heat | Not cost effective | Cost effective | Cost effective |
| Li Z., et. al. (2018) J. Chromatogr. A. 1581-82, 21-28 | For Nuclear protein-HBc. Supernatant, then 70 °C, 30 min | Heat | Not cost effective | Cost effective | Cost effective |
| Maestro B., et. al. (2008) J. Chromatogr. A. 1208, 189-196 | 15% w/w PEG 8000, 12.5% potassium phosphate, 5 min | Polymer/Salt ATPS | Not cost effective | Not cost effective | Not cost effective |
| Maestro B., et. al. (2008) J. Chromatogr. A. 1208, 189-196 | 6% w/w PEG 8000, 6% w/w dextran, 20 mM Tris pH 8, 5 min | Polymer/Salt ATPS | Not cost effective | Not cost effective | Not cost effective |
| Mao L. N., et. al. (2010) Biotechnol. Prog. 26, 1662-1670 | - | Polymer/Salt ATPS | Not cost effective | Not cost effective | Not cost effective |
| Marciniak A., et. al. (2019) Food Chem. 275, 193-196 | For alpha-lactalbumin | Pressure/pH | Not cost effective | Not cost effective | Cost effective |
| Marciniak A., et. al. (2019) Food Chem. 275, 193-196 | For beta-lactoglobulin | Pressure/pH | Not cost effective | Not cost effective | Cost effective |
| Marek W., et. al. (2013) J. Chromatogr. A. 1305, 55-63 | - | Polymer/Salt ATPS | Not cost effective | Not cost effective | Not cost effective |
| Matlschweiger A., et. al. (2017) Biotechnol. Prog. 33, 642-653 | - | Polymer | Not cost effective | Cost effective | Cost effective |
| McCabe M. S., et. al. (2008) Plant Biotechnol. J. 6, 914-929 | - | Inorganic Salt | Potential replacement | Potential replacement | Potential replacement |
| McDonald P., et. al. (2009) Biotechnol. Bioeng. 102, 1141-1151 | Protein A eluate, 0.8 mS/cm, 2.13 g PVS per g product, pH 7, 30 min | Polyelectrolyte | Not cost effective | Not cost effective | Cost effective |

| | | | | | |
|---|---|---|---|---|---|
| McDonald P., et. al. (2009) Biotechnol. Bioeng. 102, 1141-1151 | HCCF, 1.5 mS/cm, 0.57 g PVS /g Antibody, pH 7, 30 min. | Polyelectrolyte | Not cost effective | Potential replacement | Potential replacement |
| McDonald P., et. al. (2009) Biotechnol. Bioeng. 102, 1141-1151 | Protein A eluate, 0.7 mS/cm, 0.19 g PVS /g Antibody, pH 7, 30 min. | Polyelectrolyte | Not cost effective | Not cost effective | Cost effective |
| McDonald P., et. al. (2009) Biotechnol. Bioeng. 102, 1141-1151 | HCCF, 0.7 mS/cm, 0.76 g PVS /g Antibody, pH 7, 30 min. | Polyelectrolyte | Not cost effective | Not cost effective | Cost effective |
| Menkhaus T. J., et. al. (2002) 77, 148-154 | For D10-beta-glucuronidase. 30 mg PEI/g total protein | Polyelectrolyte | Potential replacement | Potential replacement | Potential replacement |
| Menkhaus T. J., et. al. (2002) 77, 148-154 | For D10-beta-glucuronidase. 100 mg PEI/g total protein | Polyelectrolyte | Not cost effective | Cost effective | Cost effective |
| Menkhaus T. J., et. al. (2002) 77, 148-154 | For D10-beta-glucuronidase. 50 mg PEI/g total protein | Polyelectrolyte | Not cost effective | Not cost effective | Cost effective |
| Menkhaus T. J., et. al. (2002) 77, 148-154 | For beta-glucuronidase. 30 mg PEI/g total protein | Polyelectrolyte | Not cost effective | Cost effective | Cost effective |
| Menkhaus T. J., et. al. (2002) 77, 148-154 | For beta-glucuronidase. 280 mg PEI/g total protein | Polyelectrolyte | Potential replacement | Potential replacement | Potential replacement |
| Menkhaus T. J., et. al. (2002) 77, 148-154 | For beta-glucuronidase. 50 mg PEI/g total protein | Polyelectrolyte | Not cost effective | Cost effective | Cost effective |
| Menkhaus T. J., et. al. (2002) 77, 148-154 | For beta-glucuronidase. 100 mg PEI/g total protein | Polyelectrolyte | Not cost effective | Cost effective | Cost effective |
| Menkhaus T. J., et. al. (2002) 77, 148-154 | For D10-beta-glucuronidase. 280 mg PEI/g total protein | Polyelectrolyte | Not cost effective | Cost effective | Cost effective |
| Menzel S., et. al. (2018) Protein Expr. Purif. 152, 122-130 | - | Heat | Potential replacement | Potential replacement | Potential replacement |
| Ng M. Y. T., et. al. (2006) 137, 134-139 | - | Heat | Not cost effective | Not cost effective | Cost effective |
| Oelmeier S.A., et. al. (2012) J. Chromatogr. A. 1252, 104-114 | For mAb 2. 16.5% w/w PEG 400, 17.45% w/w PO4, pH 6 | Polymer/Salt ATPS | Not cost effective | Not cost effective | Not cost effective |
| Oelmeier S.A., et. al. (2013) J. Chromatogr. A. 1319, 118-126 | For mAb 1. 50 mM citrate, pH 6, 15% w/w PEG 4000 | Polymer | Not cost effective | Not cost effective | Potential replacement |
| Oelmeier S.A., et. al. (2013) J. Chromatogr. A. 1319, 118-126 | For mAb 2. 50 mM citrate, pH 6, 15% w/w PEG 4000 | Polymer | Not cost effective | Not cost effective | Potential replacement |
| Oelmeier S.A., et. al. (2013) J. Chromatogr. A. 1319, 118-126 | For mAb 4. 50 mM citrate, pH 6, 15% w/w PEG 4000 | Polymer | Not cost effective | Not cost effective | Potential replacement |
| Oelmeier S.A., et. al. (2013) J. Chromatogr. A. 1319, 118-126 | For mAb 3. 21.02% w/w PEG 400, 18.99% w/w citrate, pH 6 | Polymer/Salt ATPS | Not cost effective | Not cost effective | Not cost effective |
| Ouyang J. O., et. al. (2003) Protein Expr. Purif. 32, 28-34 | - | Inorganic Salt | Not cost effective | Not cost effective | Not cost effective |
